# Nrg1/ErbB Signaling-Mediated Regulation of Fibrosis After Myocardial Infarction

**DOI:** 10.1101/2021.01.29.428912

**Authors:** Manabu Shiraishi, Atsushi Yamaguchi, Ken Suzuki

## Abstract

**RATIONALE:** Appropriate fibrotic tissue formation after myocardial infarction (MI) is crucial to maintenance of the heart’s structure. Reparative or M2-like macrophages play a vital role in post-MI fibrosis by activating cardiac fibroblasts. However, the mechanism by which post-MI cardiac fibrosis is regulated is not fully understood.

**OBJECTIVE:** We investigated the cellular and molecular mechanisms of post-MI fibrotic tissue formation, especially those related to regulation of cellular senescence and apoptosis.

**METHODS AND RESULTS:** *In vivo* and *in vitro* experiments were used to investigate the molecular and cellular mechanisms through which post-MI fibrosis occurs, with a focus on the role of M2-like macrophages. Microarray analysis revealed that CD206^+^F4/80^+^CD11b^+^ M2-like macrophages collected from mouse hearts on post-MI day 7 showed increased expression of neuregulin 1 (Nrg1). Nrg1 receptor epidermal growth factor receptor ErbB was expressed on cardiac fibroblasts in the infarct area. In cardiac fibroblasts in which hydrogen peroxide-induced senescence, M2-like macrophage-derived Nrg1 suppressed both senescence and apoptosis of the fibroblasts, whereas blockade of ErbB function significantly accelerated. M2-like macrophage-derived Nrg1/ErbB/PI3K/Akt signaling, which was shown to be related to anti-senescence, was activated in damaged cardiac fibroblasts. Interestingly, systemic blockade of ErbB function in MI model mice enhanced senescence and apoptosis of cardiac fibroblasts and exacerbated inflammation. Further, increased accumulation of M2-like macrophages resulted in excessive progression of fibrosis in post-MI murine hearts. The molecular mechanism underlying regulation of fibrotic tissue formation in the infarcted myocardium was shown in part to be attenuation of apoptosis and senescence of cardiac fibroblasts by activation of Nrg1/ErbB/PI3K/Akt signaling.

**CONCLUSIONS:** M2-like macrophage-mediated regulation of Nrg1/ErbB signaling, have a substantial effect on fibrotic tissue formation in the infarcted adult mouse heart, is critical for suppressing the progression of senescence and apoptosis of cardiac fibroblasts.

## INTRODUCTION

Myocardial infarction (MI) is a leading cause of mortality and disability worldwide. Heart failure is common among survivors of acute MI, resulting from the adverse ventricular remodeling that follows MI.^1, 2^ Because the regenerative capacity of the human heart does not fully compensate for the loss of cardiomyocytes that occurs with MI, formation of connective tissue is essential to maintenance of the integrity and rigidity of the heart. However, the mechanism by which cardiac fibrosis is regulated after MI is not fully understood.

MI results in permanent loss of hundreds of millions of cardiomyocytes.^3^ Studies have shown that in the infarct area even non-cardiomyocytes, including fibroblasts, disappear in large quantities by apoptosis and that cellular senescence-associated defects occur in cardiac repair after MI.^4, 5^ Apoptosis plays an important role in the disappearance of infiltrated immune cells and cardiac interstitial cells after MI.^5^ Because senescence and apoptosis of both cardiomyocytes and fibroblasts are deeply involved in the pathophysiology of adverse left ventricular remodeling and myocardial rupture after MI,^6^ understanding the molecular mechanisms by which senescence and apoptosis are regulated during the post-MI tissue repair process is important. Cellular senescence and apoptosis are processes of growth arrest that occur with age and in response to cellular stress and damage, and they limit proliferation of cells.^7, 8^ Senescent cells possess a complex phenotype and are characterized by cell cycle arrest mediated via p16 and p53/p21 pathways, with some ultimately manifesting a unique secretory phenotype known as the senescence-associated secretory phenotype (SASP).^9^ Cell cycle arrest plays a central role in the senescent phenotype of adult cardiomyocytes, and induction of cell cycle reentry of adult cardiomyocytes promotes post-MI cardiac repair.^10^ Administration of anti-apoptotic substances to rats after MI has been shown to suppress cardiomyocyte apoptosis, which in turn decreases infarct size and ameliorates the cardiac dysfunction that typically follows MI.^11^ Even non-cardiomyocytes in the infarct area, including fibroblasts, undergo apoptosis.^4, 5^ Therefore, senescence and/or apoptosis of cardiac fibroblasts may be involved in the tissue repair process that follows MI. Senescent cardiac fibroblasts, in which expression of the major senescence regulator p53 is significantly upregulated, have been shown to accumulate markedly in the mouse heart after MI, with the upregulation of p53 shown to limit cardiac collagen production, and conversely, the inhibition of p53 shown to increase reparative fibrosis.^1^ However, the methods used in the aforementioned studies are limited in their ability to identify the particular subset of immune cells involved in the post-MI fibrotic process as well as the inherent cellular and molecular mechanisms by which fibroblast senescence and collagen production are regulated. Whereas fibrosis after MI is crucial to maintaining myocardial structure, excessive fibrosis leads to eventual heart failure. Therefore, achieving equilibrium between profibrotic and anti-fibrotic environments is important for a good regenerative outcome. For an understanding of the fibrosis-based tissue repair process, intercellular communication between senescent and apoptotic fibroblasts and surrounding immune cells must be clarified.

Investigators have shown macrophages to be essential for regeneration of the neonatal mouse heart,^12^ and we have shown that reparative or M2-like macrophages play a pivotal role in the formation of fibrous tissue following MI by promoting proliferation and activation of cardiac fibroblasts.^13^ We hypothesized that M2-like macrophages play a vital role in attenuating apoptosis and senescence of cardiac fibroblasts, both of which are profoundly involved in the regulation of fibrotic processes after MI, and we investigated the molecular mechanism by which this particular subset of immune cells effectuates this regulation.

Neuregulin 1 (*Nrg1*) is one of the neuregulin genes (*Nrg1–Nrg4*) belonging to the family of epidermal growth factor genes,^14, 15^ and Nrg1/epidermal growth factor receptor (ErbB) signaling systems play essential roles in the protection and proliferation of cardiomyocytes in response to injury.^16–19^ Although association between Nrg1 and protection of cardiomyocytes has been studied over several decades,^16–19^ the exact mechanism through which Nrg1 protects cardiac fibroblasts and the mechanism through which Nrg1 participates in post-MI regeneration have not been elucidated. A recent study showed that Nrg1 enhances proliferation and viability of normal human ventricular cardiac fibroblasts, with the Nrg1 activity linked to Nrg1/ErbB signaling.^20^ Results of a study in a mouse model of myocardial hypertrophy indicated that Nrg1 has an anti-fibrotic effect in the heart, generated by anti-inflammatory Nrg1/ErbB signaling in macrophages.^21^ Furthermore, Nrg1-loaded poly-microparticles were used in a later study to induce macrophage polarization toward an anti-inflammatory phenotype, which prevented their transition toward the inflammatory phenotype and enhanced cardiac repair after MI.^22^ However, whether M2-like macrophages function to ameliorate senescence and apoptosis of cardiac fibroblasts through Nrg1/ErbB signaling activity *in vivo* under post-MI conditions is unknown. Although the contribution of Nrg1/ErbB signaling in macrophages and macrophage polarization toward an anti-inflammatory phenotype to cardiac tissue repair was investigated in these studies, the significance of Nrg1/ErbB signaling activity in cardiac fibroblasts with respect to the anti-fibrotic effect of Nrg1 has not been clarified. We established what we believe to be an appropriate *in vitro* model for delineation of the precise roles of M2-like macrophage-derived Nrg1 after MI in anti-senescence, anti-apoptosis, and the fibrotic phenotype of fibroblasts. Moreover, we used this model and an *in vivo* model to investigate the molecular and cellular mechanisms by which post-MI fibrosis occurs, with a focus on the role of M2-like macrophages. We aimed to shed light on the pathophysiology of fibrosis after MI and provide information that will contribute to ultimate development of new therapeutic strategies that exploit Nrg1-driven senescence and apoptosis of cardiac fibroblasts.

## METHODS

### Data Availability

The data that support the findings of this study are available from the corresponding author upon reasonable request.

Microarray data: Gene Expression Omnibus GSE69879 (https://www.ncbi.nlm.nih.gov/geo/query/acc.cgi?acc=GSE69879)

Please see the Supplemental Materials for Methods and Major Resources.

## RESULTS

### Cardiac Fibroblasts Undergo Apoptosis and Senescence After MI

We investigated cellular senescence and apoptosis in a mouse MI model, which was established by means of coronary artery ligation. Obvious fibrotic tissue formation and increased myocardial expression of fibrosis-associated genes (*αSMA* [alpha-smooth muscle actin gene], *Col1a1*, and *Col3a1*) were observed in the infarct area as soon as post-MI day 7 (Figure IA–B in the Data Supplement). These changes were associated with an increase in the number of thymocyte antigen 1 (Thy1)^+^ fibroblasts and Thy1^+^αSMA^+^ myofibroblasts in the infarct area (Figure IC–D in the Data Supplement), which peaked on post-MI day 7. Interestingly, approximately 40% of Thy1^+^ fibroblasts in the infarct area were positive for cleaved caspase 3 on post-MI day 7, and approximately 15% were positive for cleaved caspase 3 on post-MI day 28, indicative of robust apoptosis of cardiac fibroblasts (Figure 1A). Senescence-associated β-galactosidase (SA-β-gal)- positive cells were also found in the infarct area. Numerous SA-β-gal-positive cells were spindle-shaped and had many cytoplasmic processes and thus appeared to be fibroblasts (Figure 1B). Additionally, myocardial expression of senescence-associated genes (*p16*, *p2*, and *Glb1* [beta 1 galactosidase gene])^1, 7, 8, 23, 24^ and inflammation-associated genes (*CCL3*, *IL-6*, and *TNF*) was upregulated in the infarct area in comparison to that in the non-infarct (remote) area (Figure 1C and IE in the Data Supplement). Importantly, myocardial expression of reparative gene *Nrg1* was upregulated in the infarct area, peaking on post-MI day 7 (Figure 1D). Immunohistochemistry (IHC) showed that most Thy1^+^ cardiac fibroblasts that had accumulated in both the infarct and remote areas expressed ErbB2 and ErbB4, which are Nrg1 coreceptors^19, 25, 26^ (Figure 1E). Taken together, these results indicate that apoptosis and senescence of cardiac fibroblasts occur during fibrotic tissue formation in the post-MI heart and that cardiac fibroblasts express 2 Nrg1 receptors, ErbB2 and ErbB4, in response to ischemic injury.

**Figure 1.**
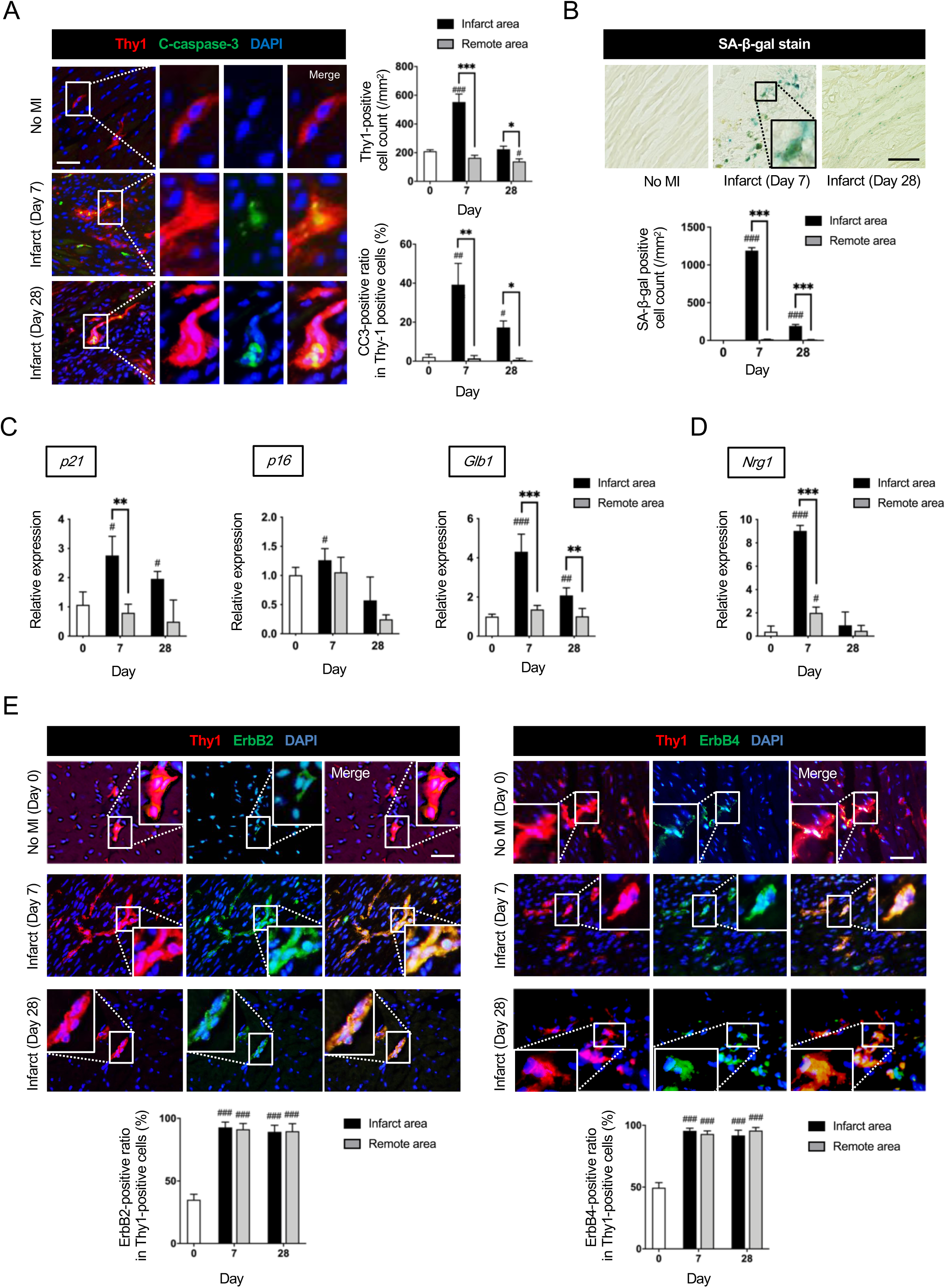
Myocardial infarction (MI) induces apoptosis and senescence of cardiac fibroblasts and upregulates Nrg1 expression in infarcted myocardium. **A,** Double immunofluorescence staining for Thy1 and cleaved caspase 3 (CC3), performed on post-MI days 7 and 28, revealed a significantly increased number of apoptotic cardiac fibroblasts in the infarct area, relative to the number seen in the remote area. Scale bars: 100 µm. *n=*4 samples each. **B**, Staining for senescence-associated beta-galactosidase (SA-β-gal) on post-MI days 7 and 28 revealed a significantly increased accumulation of spindle-shaped senescent cells in the infarct area relative to that in the remote area. Scale bars: 100 µm. **C**, Quantitative reverse transcription-polymerase chain reaction (qRT-PCR) revealed significant upregulation of senescence-associated genes (*p16* and *Glb1*) in the infarct area on post-MI day 7 or 28 or both. *n=*4 samples each. **D**, qRT-PCR analysis confirmed significant upregulation of *Nrg1* in the infarct area on post-MI day 7. *n=*4 samples each. **E**, On immunohistochemistry, the Thy1^+^ErbB2^+^ fibroblast/Thy1^+^ fibroblast ratio and the Thy1^+^ErbB4^+^ fibroblast/Thy1^+^ fibroblast ratio determined on post-MI day 7 and 28 were found to be significantly increased. Scale bars: 100 µm. *n=*4 samples each. Gene expression levels relative to those in non-MI heart (Day 0) are given. Mean±SEM values are shown \**P*<0.05, \*\**P*<0.01, \*\*\**P*<0.005 versus the remote area; ^#^*P*<0.05, ^##^*P*<0.01, ^###^*P*<0.005 versus non-MI heart; 1-way ANOVA.

### M2-Like Macrophages Accumulate in the Infarct Area and Express Nrg1

IHC showed that CD206^+^ M2-like macrophages had accumulated in the infarct area, with peak accumulation observed on post-MI day 7 (Figure 2A). Interestingly, this change in M2-like macrophages corresponded to the change that occurred in cardiac fibroblasts after MI (Figure IC in the Data Supplement). We further confirmed that CD206^+^ cells were also positive for F4/80 in both normal and infarcted hearts (Figure 2B). Microarray analysis showed that the molecular signature of CD206^+^F4/80^+^CD11b^+^ M2-like macrophages collected from infarcted hearts on post-MI day 7 differed from that of M2-like macrophages collected from normal hearts (Figure IIA in the Data Supplement). A range of anti-inflammatory and reparative genes was upregulated in M2-like macrophages in comparison to expression of the same genes in M2-like macrophages obtained from normal hearts (Figure IIB in the Data Supplement; the full dataset is available in the Gene Expression Omnibus [GEO] database; GSE69879). Importantly, gene ontology analysis showed association between M2-like macrophages obtained from infarcted hearts and regulation of apoptosis and cell death (Figure IIC in the Data Supplement). By searching HomoloGene (https://www.ncbi.nlm.nih.gov/homologene) for genes related to proliferation and viability, we focused on the increased expression level of *Nrg1* in M2-like macrophages obtained from infarcted hearts (Figure 2C), which was confirmed by quantitative reverse transcription-polymerase chain reaction (qRT-PCR) and IHC (Figure 2D–E). The data indicate that M2-like macrophages are a source of Nrg1 after MI.

**Figure 2.**
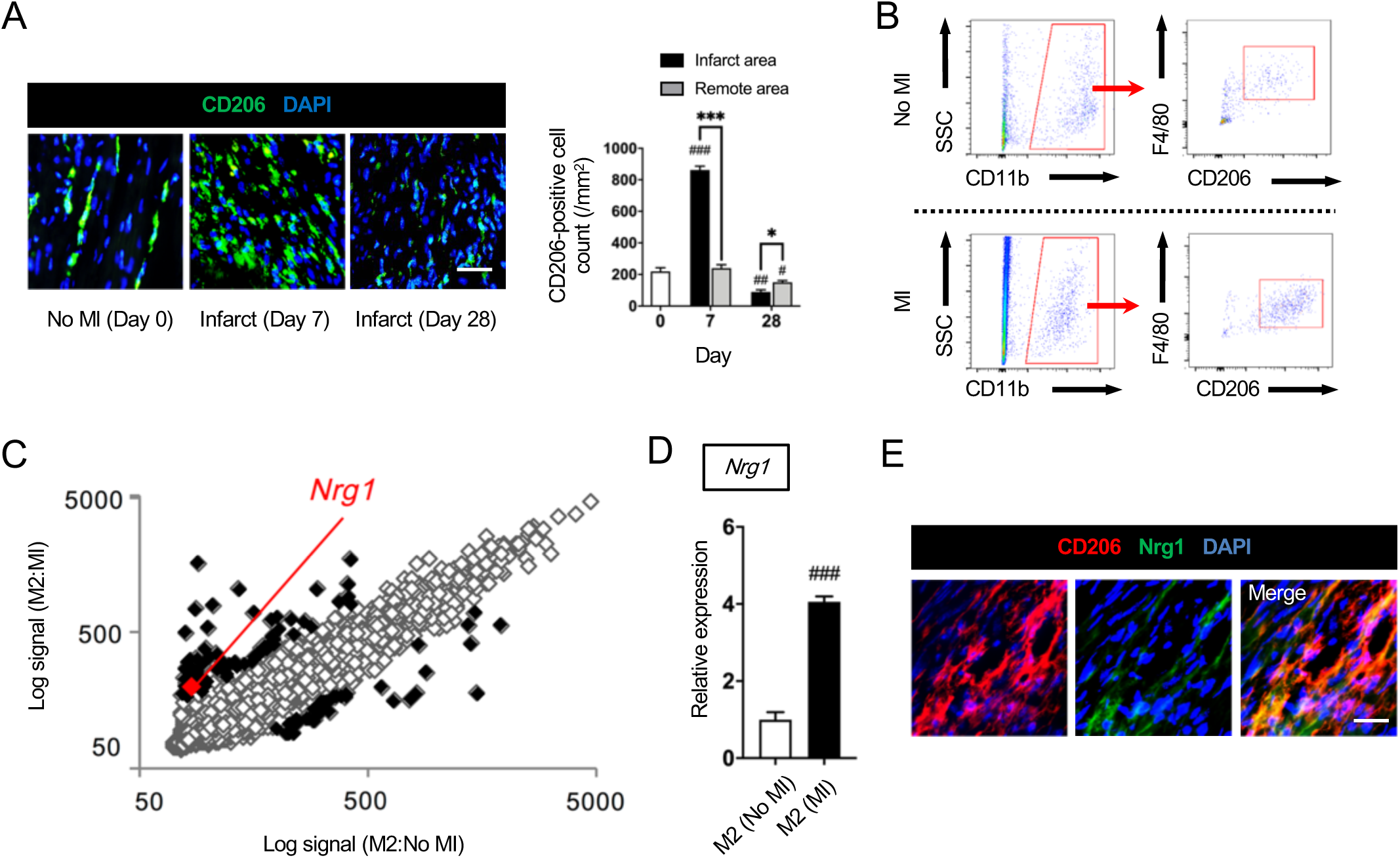
M2-like macrophages accumulate in the infarct area and express Nrg1. **A**, Immunohistochemistry revealed increased accumulation of CD206^+^ M2-like macrophages in the infarct area, which peaked on post-MI day 7. Scale bars: 100 µm. *n=*4 samples each. Mean±SEM values are shown. \**P*<0.05, \*\**P*<0.01, \*\*\**P*<0.005 versus the remote area; ^#^*P*<0.05, ^##^*P*<0.01, ^###^*P*<0.005 versus non-MI heart; 1-way ANOVA. **B**, Flow cytometric analysis confirmed the presence of CD206^+^F4/80^+^CD11b^+^ M2-like macrophages in normal (non-MI) hearts of adult C57BL/6 mice and in hearts of mice subjected to MI. *n=*6 samples each. **C**, Microarray analysis showed the M2 macrophages after MI to have a different expression profile from that of M2 macrophages in hearts not subjected to MI. The scatter plot shows upregulation of 70 genes and downregulated of 39 genes in M2-like macrophages after MI compared to M2-like macrophages. Among the upregulated genes is neuregulin 1 (*Nrg1*). **D**, Quantitative reverse transcription-polymerase chain reaction confirmed post-MI upregulation of *Nrg1* in CD206^+^F4/80^+^CD11b^+^ cardiac M2-like macrophages after MI. *n=*4 samples each. Gene expression levels relative to those in M2-like macrophages from normal hearts are shown. Mean±SEM values are shown. ^#^*P*<0.05, ^##^*P*<0.01, ^###^*P*<0.005 versus M2-like macrophages from normal hearts, 2-tailed, unpaired Student’s *t-*test. **E**, Double immunofluorescence staining revealed Nrg1 expression on the surface of CD206^+^ M2-like macrophages. Scale bars: 100 µm. *n=*4 samples each.

### Bone Marrow-Derived Macrophages Attenuate Hydrogen Peroxide (H_2_O_2_)-Induced Apoptosis and Senescence of Cardiac Fibroblasts Via Nrg1 Secretion

We investigated the role of macrophages in regulating senescence and apoptosis of fibroblasts in an *in vitro* coculture model using a Boyden Chamber system in which cells can be independently cultured or genetically analyzed without being mixed together^13, 27^ (Figure IIIA in the Data Supplement). Hydrogen peroxide (H_2_O_2_) was used to induce senescence of fibroblasts (Figure IIIB–C in the Data Supplement). Bone marrow-derived macrophages (BMDMs) that were cocultured with the senescent fibroblasts were of an M2-like macrophage phenotype (Figure IVA–B in the Data Supplement). As observed *in vivo* after MI (Figure 2C–E and Figure 1E), increased expression of *Nrg1* was seen in BMDMs (Figure VA in the Data Supplement) and increased expression of *ErbB2* and *ErbB4* was seen in cardiac fibroblasts (Figure VB–C in the Data Supplement) 48 hours after the start of coculture. Phase-contrast microscopy (Figure 3A) showed that fibroblasts treated with H_2_O_2_ had an enlarged, flattened, senescent morphology, but after coculture with BMDMs the fibroblasts took on a healthy, spindle-shaped form. Importantly, addition of anti-ErbB antibody (Ab), which is a competitive inhibitor of Nrg1, changed the gross morphology of fibroblasts such that they resembled the fibroblasts treated with H_2_O_2_. Treatment with recombinant Nrg1 restored the senescent fibroblasts to a healthy spindle-shaped form. Approximately 20% of fibroblasts treated with H_2_O_2_ were found to be positive for senescence-associated β-galactosidase (SA-β-gal) (Figure 3B). This change was significantly attenuated by coculture with BMDMs, whereas addition of anti-ErbB Ab eliminated the anti-senescence effect of the BMDMs. Furthermore, treatment with recombinant Nrg1 suppressed senescence. Expression of apoptosis-related cleaved caspase 3 (Figure 3C) was similar to that of SA-β-gal. Conversely, expression of proliferation-related Ki-67 was significantly attenuated by H_2_O_2_ treatment and restored by co-culture with BMDMs. Effects of the BMDMs were diminished by anti-ErbB Ab, whereas recombinant Nrg1 improved H_2_O_2_-induced functional deterioration. Together, these results show that Nrg1 derived from BMDMs reduces both apoptosis and senescence of cultured fibroblasts and promotes their proliferation.

**Figure 3.**
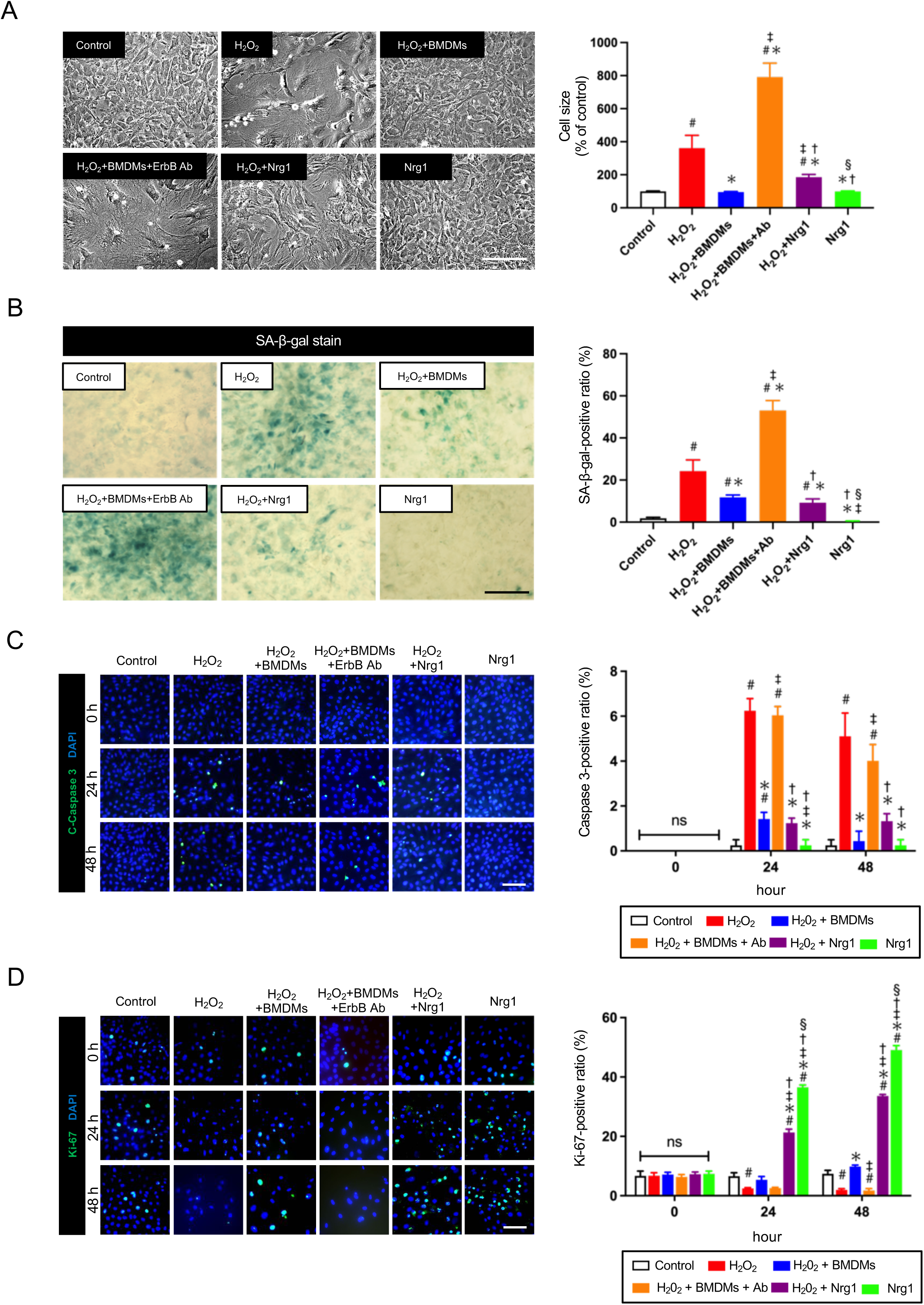
Bone marrow-derived macrophages (BMDMs) attenuate H_2_O_2_-induced apoptosis and senescence of cardiac fibroblasts via Nrg1 secretion. **A,** Representative phase-contrast microscopy images. Treatment with a hydrogen peroxide (H_2_O_2_) solution changed the spindle-shaped appearance of cardiac fibroblasts to a significantly enlarged, flattened morphology. Addition of bone marrow-derived macrophages (BMDMs) altered the senescent morphology to a normal, spindle-shaped morphology. After addition of anti-ErbB antibody (Ab), fibroblasts displayed the same gross morphology as that of senescent fibroblasts treated with H_2_O_2_. Likewise, addition of recombinant neuregulin 1 (Nrg1) changed the gross morphology to a spindle shape. Scale bars: 100 µm. *n=*4 samples each. **B.** Staining for SA-β-gal showed that senescence was exacerbated in H_2_O_2_-treated fibroblasts but not in fibroblasts cocultured with BMDMs. This suppression of senescence was attenuated by coculture with the anti-ErbB Ab. Nrg1 suppressed fibroblast senescence. Scale bars: 100 µm. *n=*4 samples each. **C.** Apoptosis of cardiac fibroblasts (ratio of cleaved caspase 3^+^DAPI^+^ fibroblasts to DAPI^+^ fibroblasts) was increased in coculture with H_2_O_2_ but decreased in coculture with BMDMs. This decrease in apoptosis was eliminated by the anti-ErbB Ab. Nrg1 suppressed apoptosis. Nuclei were counterstained with 4ʹ,6-diamidino-2-phenylindole (DAPI). *n=*4 samples each. Scale bars: 100 µm. **D**, Proliferation of cardiac fibroblasts (ratio of Ki-67^+^ and DAPI^+^ fibroblasts to DAPI^+^ fibroblasts) was decreased under H_2_O_2_ treatment but increased when cells were cocultured with BMDMs. This enhanced proliferation was suppressed by the anti-ErbB Ab. Nrg1 accelerated proliferation. Nuclei were counterstained with DAPI. Scale bars: 50 µm. *n=*4 samples each. Mean±SEM values are shown. ^#^*P*<0.05 versus normal cardiac fibroblasts (control), \**P*<0.05 versus H_2_O_2_, ^‡^*P*<0.05 versus H_2_O_2_+BMDMs, ^†^*P<*0.05 versus H_2_O_2_+BMDMs+Ab, ^§^*P*<0.05 versus H_2_O_2_+Nrg1; 1-way ANOVA.

### BMDMs Promote Activation of Fibroblasts and Collagen Synthesis

Immunocytological staining showed that, although treatment with H_2_O_2_ alone did not affect conversion of cardiac fibroblasts into αSMA^+^ myofibroblasts, the phenotypic change progressed in coculture of cardiac fibroblasts with BMDMs. Furthermore, addition of anti-ErbB Ab enhanced this BMDM-induced activation of fibroblasts. Addition of recombinant Nrg1 did not affect conversion of cardiac fibroblasts (Figure 4A). Changes in synthesis of types I and III collagen in fibroblasts in response to H_2_O_2_, BMDMs, and recombinant Nrg1 were similar to changes in αSMA expression (Figure 4B–C). These results suggest that BMDMs, which are of an M2-like phenotype (Figure IVA–B in the Data Supplement), induce activation of fibroblasts for conversion to myofibroblasts and progression of collagen synthesis. The important take-away point to be drawn from these results is that that cellular senescence and apoptosis alone do not induce activation of fibroblasts for conversion to myofibroblasts and progression of collagen synthesis; the presence of BMDMs is required. Osteopontin (secreted phosphoprotein 1 [SPP1]) is a major mediator of M2-like macrophage-induced cardiac fibroblast activation.^13^ *Spp1* expression was found to be increased in BMDMs in response to H_2_O_2_ (Figure VIA in the Data Supplement). Conversely, other profibrotic factors, including transforming growth factor beta and platelet derived growth factor subunit A,^28, 29^ were found not to be upregulated in M2-like macrophages (Figure VIB in the Data Supplement).

**Figure 4.**
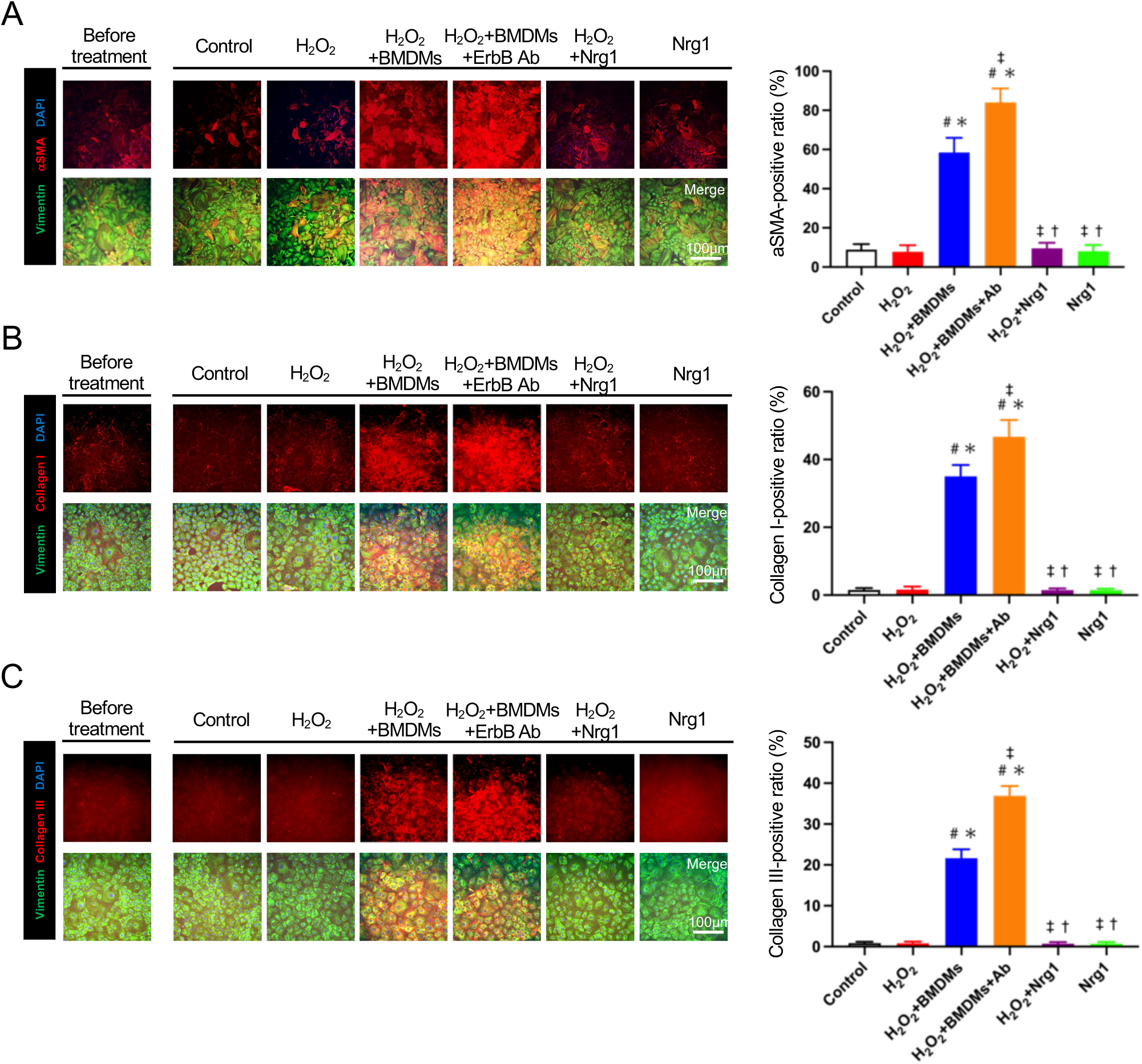
Bone marrow-derived macrophages (BMDMs) promote fibroblast activation and collagen synthesis. **A–C**. Representative images of cardiac fibroblasts stained immunocytochemically for (A) vimentin and αSMA, (B) vimentin and collagen I, and (C) vimentin and collagen III. The staining was performed 48 hours after the start of culture. **A,** Activation of cardiac fibroblasts (ratio of vimentin^+^ and αSMA^+^ myofibroblasts to vimentin^+^ fibroblasts) was equal under treatment with hydrogen peroxide (H_2_O_2_) but markedly increased when cells were cocultured with bone marrow-derived macrophages (BMDMs). This increase in activation was accelerated by the anti-ErbB antibody (Ab). Addition of recombinant neuregulin 1 (Nrg1) did not affect activation. Scale bars: 100 µm. *n=*4 samples each. **B.** Collagen I synthesis was equal under treatment with H_2_O_2_ but significantly increased when cells were cocultured with BMDMs. This increase in production was enhanced by treatment with anti-ErbB Ab. Addition of Nrg1 did not affect collagen I synthesis. Scale bars: 100 µm. *n=*4 samples each. **C.** Collagen III synthesis was similar to collagen I synthesis. Scale bars: 100 µm. *n=*4 samples each. Mean±SEM values are shown. ^#^*P*<0.05 versus normal cardiac fibroblasts (control), \**P*<0.05 versus H_2_O_2_, ^‡^*P*<0.05 versus H_2_O_2_+BMDMs, ^†^*P*<0.05 versus H_2_O_2_+BMDMs+Ab, ^§^*P*<0.05 versus H_2_O_2_+Nrg1; 1-way ANOVA.

### Phosphatidylinositol 3-Kinase/Protein Kinase B Signaling is Associated with BMDM-Attenuated Apoptosis and Senescence of Cardiac Fibroblasts Through Nrg1

We next investigated the molecular mechanism underlying attenuation of apoptosis and senescence of cardiac fibroblasts by BMDM-derived Nrg1 in an *in vitro* setting. The phosphatidylinositol 3-kinase (PI3K)/protein kinase B (Akt) signaling pathway is downstream of the ErbB pathway. Western blot analysis showed that H_2_O_2_ treatment suppressed PI3K/Akt activation in fibroblasts (Figure 5A). Coculture with BMDMs and treatment with recombinant Nrg1upregulated the signaling activity, whereas addition of anti-ErbB Ab suppressed the signaling activity. To determine the possible relation between senescence-associated p21/p16 and Nrg1/ErbB/PI3K/Akt signaling pathways, we examined mRNA expression levels of senescence-associated genes (*p21*, *p16*, and *Glb1)* (Figure 5B), cell cycle-associated genes (cyclin-dependent kinase 4 [*Cdk4*], cyclin-dependent kinase 6 [*Cdk6*], cyclin-dependent kinase 2 [*Cdk2*], and *Ki-67*) (Figure 5C), a p53 suppressor gene (murine double minute 2 [*MDM2*]) (Figure 5D), a cell survival-associated gene (mechanistic target of rapamycin [*mTOR*]) (Figure 5E), and an SASP-associated gene (interleukin-6 [*IL-6*]) (Figure 5F). Expression levels of senescence-associated genes (*p16* and *p21*) and the SASP-associated gene (*IL-6*) were significantly higher and expression levels of cell cycle-associated genes (*Cdk4*, *Cdk6*, *Cdk2*, and *Ki-67*) were markedly lower in fibroblasts treated with H_2_O_2_ than in non-treated (control) fibroblasts. Importantly, these changes in expression were reversed by coculture with BMDMs. Anti-ErbB Ab exacerbated senescence and cell cycle arrest, whereas recombinant Nrg1 suppressed senescence and enhanced proliferation. BMDMs enhanced or restored expression of p53 suppressor-related *MDM2* and cell survival-related *mTOR*. These results indicate that the PI3K/Akt signaling pathway activation in cardiac fibroblasts operates downstream of the BMDM-derived Nrg1/ErbB system and has a suppressive effect on cell cycle arrest and senescence. This signaling activity is also likely to increase cell proliferation and survival (Figure 5G).

**Figure 5.**
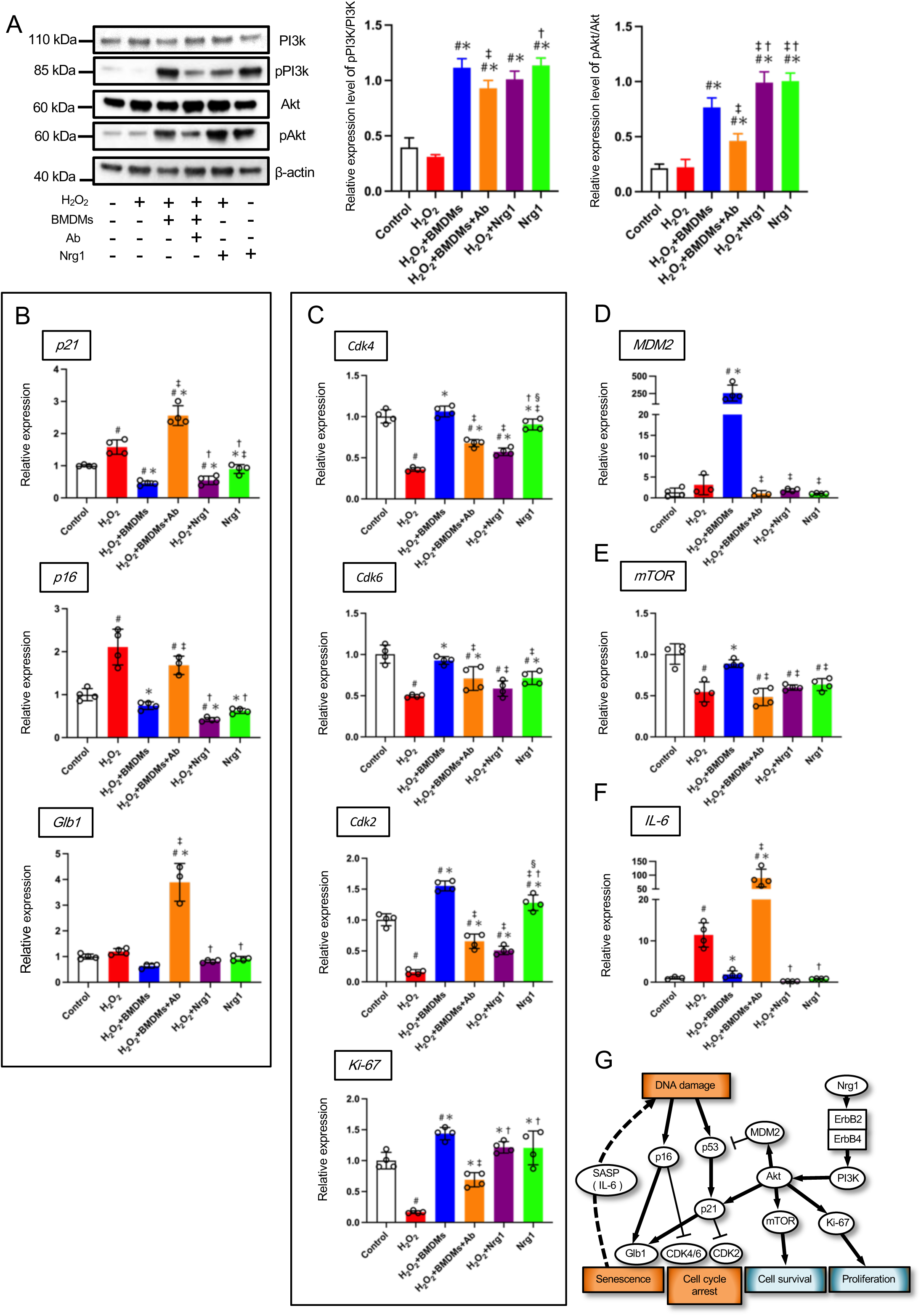
PI3K/Akt signaling pathway is associated with BMDM-attenuated apoptosis and senescence of cardiac fibroblasts through Nrg1. **A.** Representative bands of PI3K, pPI3K, Akt, pAkt, and β-actin in cardiac fibroblasts treated (or not treated) with hydrogen peroxide (H_2_O_2_), in those cocultured with bone marrow-derived macrophages (BMDMs), and those treated (or not treated) with anti-ErbB antibody (Ab)/recombinant neuregulin 1 (Nrg1). Treatment and coculture lasted 48 hours. Bar graph shows quantification of relative pPI3k/PI3k and pAkt/Akt. H_2_O_2_ alone did not affect activation of PI3K/AKT signaling in fibroblasts. Addition of BMDMs significantly activated the signaling pathway and, conversely, addition of the anti-ErbB Ab impaired it. Nrg1 re-stimulated the signaling pathway. *n*=4 samples each. **B.** Quantitative reverse transcription-polymerase chain reaction analysis of cardiac fibroblasts having been treated (or not) with H_2_O_2_, co-cultured with BMDMs, and treated (or not) with anti-ErbB Ab/Nrg1. Treatment and coculture lasted 48 hours. Expression levels of senescence-associated genes (*p21, p16*, and *Glb1*) were increased in cells treated with H_2_O_2_ but decreased with the addition of BMDMs. Addition of the anti-ErbB Ab exacerbated senescence and, conversely, addition of Nrg1 suppressed it. *n*=4 samples each. **C.** Expression of cell cycle-associated genes (*Cdk4*, *Cdk6*, *Cdk2*, and *Ki-67*) were suppressed in cells treated with H_2_O_2_ but restored in cells cocultured with BMDMs. Addition of anti-ErbB Ab decreased such expression and, conversely, addition of Nrg1 restored such expression. *n*=4 samples each. **D**, Expression of the p53 suppressor gene (*MDM2*) was significantly increased in cells cocultured with BMDMs. *n*=4 samples each. **E**, Expression of cell survival-associated gene (*mTOR*) was suppressed in cells treated with H_2_O_2_ but restored in cells cocultured with BMDMs. Addition of anti-ErbB Ab re-suppressed the expression and, conversely, addition of Nrg1 restored the *mTOR* expression. **F**, Expression of senescence-associated secretory phenotype-associated gene (*IL-6*) was significantly increased in cells treated with H_2_O_2_ and in those cocultured with BMDMs, with ErbB Ab having been added. *n*=4 samples each. **G**, Schematic representation and overview of the Nrg1/PI3K/AKT pathway. H_2_O_2_ treatment contributes to the development of cellular damage, which leads to senescence, cell cycle arrest. Nrg1 binding to coreceptor ErbB2/ErbB4 leads to activation of PI3K/AKT and inactivation of p53 and p21, which results in anti-senescence, cell survival and proliferation. Arrowheads indicate stimulation, whereas hammerheads represent inhibition. Gene expression levels relative to the normal cardiac fibroblasts (control) are given. Mean±SEM values are shown. ^#^*P*<0.05 versus control, \**P*<0.05 versus H_2_O_2_, ^‡^*P*<0.05 versus H_2_O_2_+BMDMs, ^†^*P*<0.05 versus H_2_O_2_+BMDMs+Ab, ^§^*P*<0.05 versus H_2_O_2_+Nrg1; 1-way ANOVA.

### In Vivo Inhibition of Nrg1 Signaling Exacerbates Fibrosis

We used trastuzumab to clarify the role of Nrg1 in suppressing post-MI senescence and apoptosis of cardiac fibroblasts *in vivo*. Trastuzumab is an anti-ErbB type 2 (ErbB2) monoclonal Ab that binds to the extracellular juxtamembrane domain of ErbB2.^30, 31^ On the basis of the *in vitro* coculture model, we hypothesized that trastuzumab administration would eliminate the anti-apoptotic and anti-senescence effects of Nrg1 and therefore allow senescence of cardiac fibroblasts to proceed, which would result in excessive fibrosis in the infarcted myocardium. Intraperitoneal trastuzumab injections did not affect mRNA expression levels of genes associated with senescence, the SASP, or activation of fibroblasts to myofibroblasts in the normal mouse heart (Figure VIIA–C in the Data Supplement). We induced MI in mice surgically and then injected either trastuzumab or vehicle intraperitoneally (Figure VIIIA in the Data Supplement). Trastuzumab did not affect post-MI mortality or body weight (Figure VIIIB–C in the Data Supplement). Gene expression profiles in cardiac fibroblasts resident in infarcted myocardium suggested that senescence was augmented by trastuzumab administration in the infarct area (Figure 6A). The changes in gene expression corresponded to increases in the number of senescent cardiac fibroblasts and cardiac cells in the infarct area (Figure 6B–C). Additionally, trastuzumab administration increased the proportion of fibrotic tissue in the infarct area (Figure 7A). In fact, the numbers of Thy-1^+^ fibroblasts and αSMA^+^Thy1^+^ myofibroblasts were increased in association with upregulation of *αSMA*, *Col1a1*, and *Col3a*1 in the infarct area (Figure 7B–C). Considering these *in vitro* experimental results, increased inflammation (Figure 8A) and boosted accumulation of M2-like macrophages (Figure 8B) in this area by ErbB2 blockade may explain the increased fibroblast activation and augmented fibrosis. Interestingly, trastuzumab administration induced expression of apoptosis- and senescence-associated genes even in the remote area (Figure IXA–C in the Data Supplement) where there was little or no direct ischemic damage after the MI. This corresponded to the increased inflammation (Figure XA in the Data Supplement) and the augmented accumulation of both M2-like macrophages and cardiac fibroblasts in the remote myocardium (Figure XB and Figure XIA in the Data Supplement). Conversion of fibroblasts to myofibroblasts and expression of fibrosis-associated genes were also induced (Figure XI–B in the Data Supplement). Consequently, acute exacerbation of fibrosis occurred even in the remote area (Figure XII in the Data Supplement). Thus, administration of trastuzumab augmented apoptosis and senescence of cardiac fibroblasts following MI, resulting in excessive fibrosis, even in the remote non-infarct area.

**Figure 6.**
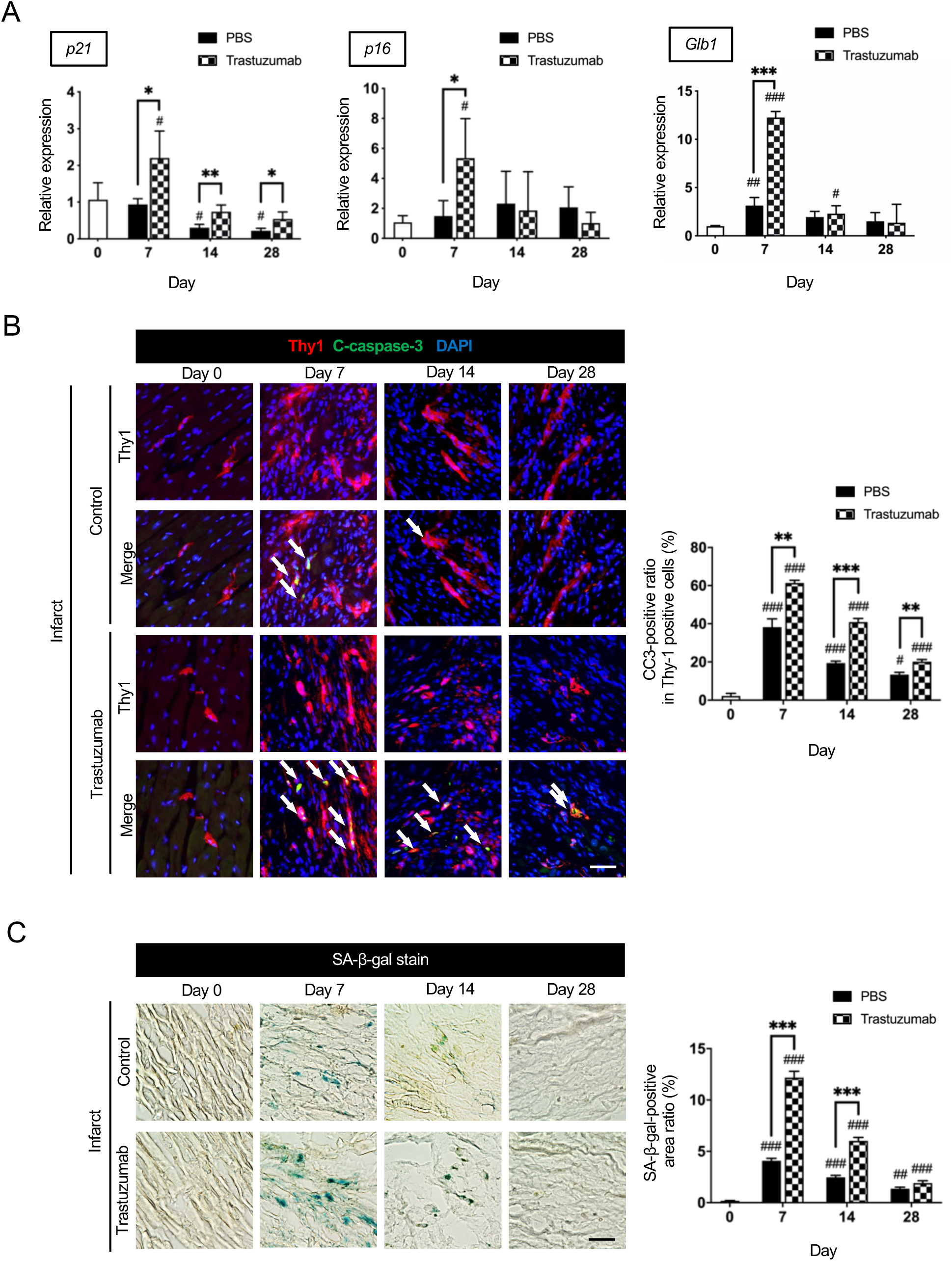
In vivo inhibition of Nrg1 signaling promotes apoptosis and senescence of cardiac fibroblasts. **A,** Quantitative reverse transcription-polymerase chain reaction (qRT-PCR) analysis showed post-myocardial infarction (MI) upregulation of senescence-associated genes (*p21*, *p16,* and *Glb1*) in the infarct area after intraperitoneal injection of the mice with trastuzumab in comparison to expression of these genes in vehicle injected post-MI heart. *n=*4 samples each. **B**, Double immunofluorescence staining of Thy1 and cleaved caspase 3 (CC3) showed the ratio of apoptotic cardiac fibroblasts in the infarct area to be increased on post-MI days 7, 14, and 28 in mice injected intraperitoneally with trastuzumab compared with that in mice injected with vehicle only. Arrow shows Thy1^+^CC3^+^ cells. Scale bars: 100 µm. *n=*4 samples each. **C**, SA-β-gal staining revealed increased senescence of cardiac cells in the infarct area on post-MI days 7 and 14 after intraperitoneal trastuzumab injection compared with senescence after vehicle injection. Scale bars: 100 µm. *n=*4 samples each. Gene expression levels relative to levels in non-MI heart (day 0) are given. Mean±SEM values are shown. \**P*<0.05, \*\**P*<0.01, \*\*\**P*<0.005 versus other sample(s); ^#^*P*<0.05, ^##^*P*<0.01, ^###^*P*<0.005 versus non-MI heart; 1-way ANOVA.

**Figure 7.**
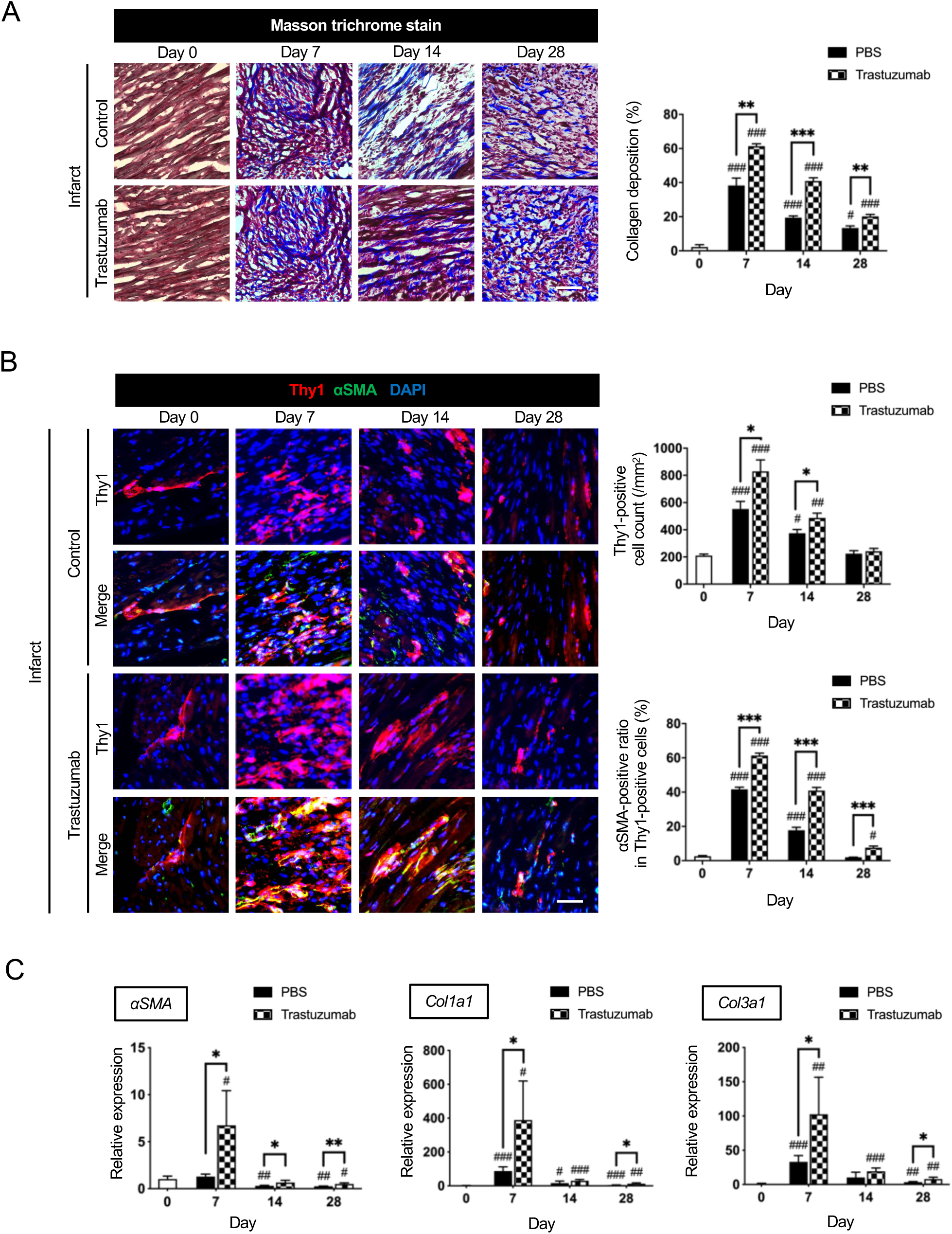
In vivo inhibition of Nrg1 signaling activates cardiac fibroblasts and exacerbates fibrosis. **A,** Masson trichrome staining demonstrated that deposition of collagen fibrils was increased in the infarct area with a time lapse. Intraperitoneal trastuzumab injection significantly increased collagen fibrils in the infarct area. Scale bars: 100 µm. *n=*4 samples each. **B**, Double immunofluorescence staining of Thy1 and αSMA showed increased accumulation and activation of cardiac fibroblasts in the infarct area following trastuzumab injection versus vehicle injection. Scale bars: 100 µm. *n=*4 samples each. **C**, Quantitative reverse transcription-polymerase chain reaction (qRT-PCR) analysis showed post-MI upregulation of fibrosis-associated genes (*αSMA*, *Col1a1*, and *Col3a1*) in mice after intraperitoneal trastuzumab injection compared with expression in mice after vehicle injection. *n=*4 samples each. Gene expression levels relative to levels in non-MI heart (day 0) are given. Mean±SEM values are shown. \**P*<0.05, \*\**P*<0.01, \*\*\**P*<0.005 versus other sample(s); ^#^*P*<0.05, ^##^*P*<0.01, ^###^*P*<0.005 versus the non-MI heart; 1-way ANOVA.

**Figure 8.**
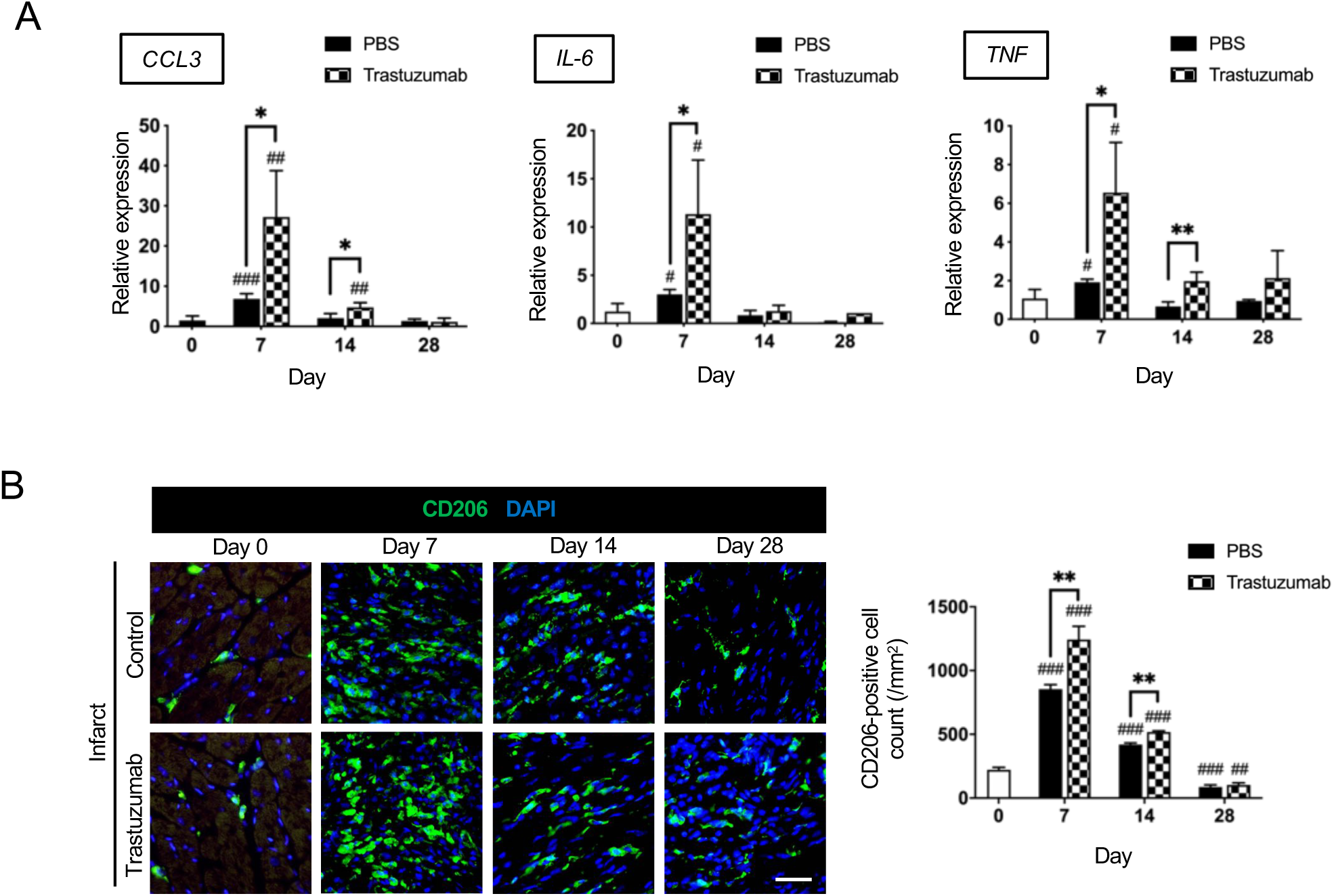
In vivo inhibition of Nrg1 signaling exacerbates myocardial inflammation and promotes accumulation of M2-like macrophages. **A,** Quantitative reverse transcription-polymerase chain reaction analysis (qRT-PCR) showed post-myocardial infarction (MI) upregulation of senescence-associated secretory phenotype-associated genes (*CCL3*, *IL-6*, and *TNF*) in mice given trastuzumab by intraperitoneal injection compared with expression in vehicle-injected mice. *n=*4 samples each. **B**, Immunohistochemistry showed intraperitoneal trastuzumab injection significantly accelerated accumulation of CD206^+^ M2-like macrophages in the infarct area with a post-MI peak on day 7. Scale bars: 100 µm. *n=*4 samples each. Gene expression levels relative to levels in non-MI heart (day 0) are given. Mean±SEM values are shown. \**P*<0.05, \*\**P*<0.01, \*\*\**P*<0.005 versus other sample(s); ^#^*P*<0.05 ^##^*P*<0.01, ^###^*P*<0.005 versus non-MI heart; 1-way ANOVA.

## DISCUSSION

Despite the clinical importance of cellular and molecular processes underlying post-MI fibrosis, the processes are not well understood. To precisely determine the roles of cardiac M2-like macrophages in apoptosis and senescence of fibroblasts, we established an *in vitro* model in which senescent cardiac fibroblasts^32^ were cocultured with BMDMs. This *in vitro* coculture model reflects the *in vivo* post-MI conditions because, under pathological conditions, senescent cells attract macrophages.^33^ Expression of both *Cdk4/6* and *Cdk2* should be stable or increased in senescent cells, but expression levels were decreased in the cardiac fibroblasts in which senescence was induced by H_2_O_2_. The mechanism is expected to involve increased expression of *p16* and *p21*, which downregulate *Cdk4/6* and *Cdk2* expression, respectively.^8, 34, 35^ Conversely, the addition of BMDMs decreased *p16* and *p21* expression and promoted the cell cycle (as evidenced by increased expression of *Cdk 4/6* and *Cdk2*). The hypertrophic response to Nrg1 is mainly dependent on Ras, whereas the anti-apoptotic and cell proliferation effects are likely dependent on Akt.^36, 37^ Therefore, Nrg1 may attenuate expression of senescence-associated genes *p21*, *p16*, and *Glb1* through PI3K/Akt pathways (Figure 5G). We observed a transient increase in *p21*, *p16*, and *Glb1* expression in the infarcted myocardium and in cultured fibroblasts. Stress-induced *p53* expression increases *p21* expression in response to DNA damage and induces reversible proliferative arrest that provides time for DNA repair and facilitates survival of cells.^38, 39^ p21, which binds to and inhibits CDK2, is important for initiation of senescence in some settings, but *p21* expression does not persist in senescent cells.^8, 34, 40, 41^ Increased *p16* expression is found in infarcted tissue. Irreversible proliferative arrest can be induced by p16, which inhibits CDK4 and CDK6.^8, 35^ Therefore, the change in expression levels of these genes (i.e., *p21* and *p16*) in both the *in vivo* and *in vitro* experiments suggests that Nrg1 is a crucial factor that controls reversible and irreversible senescence of cardiac cells after MI. Another possible mechanism for the temporal change in *p21* and *p16* expression in the *in vivo* experiment is as follows: There could be multiple types of *p21-* and *p16*-positive cells. Therefore, whereas some *p21-* and *p16*-positive cell populations disappear, other *p21-* and *p16*-positive cells may remain. We showed a possible mechanism linking M2-like macrophages and cardiac fibroblasts, which contributes to various processes that are critical to many aspects of cellular function, including cell growth and survival.^42^ Together, results of our *in vivo* and *in vitro* experiments suggest that macrophage-derived Nrg1 suppresses senescence and promotes proliferation of cardiac fibroblasts after myocardial injury by activating the ErbB/PI3K/Akt signaling pathway.

ErbB is expressed not only on the surface of fibroblasts but also on the surface of macrophages, and myeloid-specific *ErbB* deletion exacerbates myocardial fibrosis.^21^ Furthermore, Nrg1-induced macrophage polarization from an inflammatory phenotype toward an anti-inflammatory phenotype enhances cardiac repair after MI.^22^ Therefore, the question was raised whether ErbB signaling in BMDMs was simultaneously affected when anti-ErbB Ab was added to the culture medium in our coculture experiments. The exacerbated phenotypic changes characterizing senescence and apoptosis of fibroblasts might not be due to inhibition of Nrg1/ErbB signaling in the fibroblasts. However, we showed that the phenotypic changes and expression of genes denoting the senescence and apoptosis of H_2_O_2_-treated cells, with the Nrg1/ErbB signaling-induced activation of BMDMs having been excluded, were similar to those in the H_2_O_2_+BMDM+anti-ErbB Ab treated cells (Figure 3A–C and Figure 5B–C). Therefore, our data suggest that Nrg1/ErbB signaling activity has a greater anti-senescence and anti-apoptotic effect in fibroblasts than does the signaling activity related to macrophage polarization.

Nrg1 is a cytokine that belongs to a family of proteins structurally related to epidermal growth factor, and it plays essential roles in protection and proliferation of cardiomyocytes in response to injury.^16–19^ Injecting Nrg1 into adult mice induces cardiomyocyte cell cycle activity and promotes myocardial regeneration, leading to improved function after MI.^16, 18^ Nrg1 also significantly decreases apoptosis of adult cardiomyocytes under hypoxia-reoxygenation conditions.^17^ However, its function in cardiac fibroblasts in this context has not been well studied. We present new *in vivo* and *in vitro* evidence suggesting that M2-like macrophages play a vital role in attenuating senescence and apoptosis of fibroblasts through Nrg1/ErbB/PI3K/Akt signaling pathways. This inherent reparative function allows senescent cardiac fibroblasts to recover to a certain degree. Conversely, incomplete rescue of fibroblasts from senescence might lead to undesired fibrosis.

Previous studies have shown that trastuzumab efficiently stops or slows the growth of ErbB2^+^ cells *in vitro* and inhibits the ability of cells to repair damaged DNA.^30, 31^ As inflammation worsens after MI, M2-like macrophages from bone marrow accumulate at the site of damaged tissue.^43^ Our study showed that progression of senescence and apoptosis in cardiac fibroblasts and exacerbation of inflammation induced by trastuzumab increased the accumulation of M2- like macrophages, which in turn promoted activation of fibroblasts and excessive fibrosis. These results are consistent with our previous report indicating that interleukin 4-mediated M2-like macrophage activation induces conversion of fibroblasts into myofibroblasts for progression of fibrosis.^13^ Although we analyzed mRNA expression levels in sections of heart tissue and not in single cells, increased expression levels of senescence-associated genes in the trastuzumab-treated samples were considered to reflect progression of senescence. These results corresponded to results obtained *in vitro* with use of anti-ErbB Ab. We administered trastuzumab systemically, which has unclarified cardiotoxicity, to explore ErbB signaling in cardiac fibroblasts. Because cardiotoxicity of trastuzumab is non-specific, our study was limited by the fact that the ErbB receptor block achieved with anti-ErbB Ab is not specific to cardiac fibroblasts. Moreover, M2- like macrophages are not the only source of Nrg1 after MI. Apart from the M2-like macrophages we studied, endothelial cells have been identified as a potential cell type for neuregulin production and are responsible for the healing response after myocardial injury.^16–19^ Therefore, further mechanistic studies are needed to clarify the signaling pathways downstream of these factors and the clear role of Nrg1, i.e., studies that incorporate cardiac fibroblast-specific conditional *ErbB2/ErbB4*-knockout mice and macrophage-specific conditional *Nrg1*-knockout mice.

Interestingly, senescence and SASP-associated gene expression peaked a little later in the remote area than in the infarct area. One possible pathological mechanism of the remote non-infarct area is assumed to be indirect damage via SASP rather than direct cytotoxicity due to ischemia. Senescent cells autonomously induce senescence-like gene expression in surrounding non-senescent cells through the SASP.^44^ Increased release of SASP-associated substances in the infarct area might not simply induce senescence and apoptosis of cells in the infarct area; it might also have a harmful influence on non-senescent cells in the remote non-infarct area.

Although this study focused on Nrg1-induced anti-apoptosis and anti-senescence of cardiac fibroblasts, M2-like macrophages are likely to mediate supplementary benefits for cardiac repair after MI. These benefits may include reduced inflammation, activation of fibroblasts, and neovascularization, as shown in our previous study.^13^ To develop potential therapies mediated by M2-like macrophages, studies are needed to determine how the gene that encodes Nrg1 is switched on by MI and to identify other molecules that regulate apoptosis, senescence, and proliferation of cardiac fibroblasts.

In conclusion, our data provide evidence that the Nrg1/ErbB/PI3K/Akt signaling system regulates senescence and apoptosis of cardiac fibroblasts in the infarcted adult murine heart. This process might play a vital role in repair of the infarcted myocardium by regulating collagen synthesis. Therefore, this tissue repair mechanism controls the degree of rigidity and contraction of the infarcted heart, thereby determining the prognosis of MI. Better understanding of the molecular mechanism at play in the healing process of MI and subsequent remodeling may ultimately lead to new treatments. Targeted activation of M2-like macrophages might enhance the endogenous repair mechanism in senescent cardiac fibroblasts and serve as an approach to treatment for MI.

## Acknowledgments

MS and KS conceived and designed the study. MS conducted most of the experiments and acquired the data with support from AY and KS. All authors participated in the analysis and interpretation of the data. MS and KS primarily wrote and edited the manuscript with input from all other authors. We thank Ellen Knapp, PhD, and Mitchell Arico from Edanz Group (https://en-author-services.edanz.com/ac) for editing a draft of this manuscript. We wish to thank Ms. Wendy Alexander-Adams for her assistance in reporting our findings in English. This project was supported by the Uehara Memorial Foundation, SENSHIN Medical Research Foundation, a Grant-in-Aid for Scientific Research (C), the Mochida Memorial Foundation for Medical and Pharmaceutical Research, Takeda Science Foundation, and Public Trust Surgery Research Fund.

## SUPPLEMENTAL MATERIALS

Extended Methods Online Figures I–XII Online Table I–III

## Nonstandard Abbreviations and Acronyms

Akt: protein kinase B
αSMA: alpha-smooth muscle actin
BMDM: bone marrow-derived macrophage
Cdk: cyclin-dependent kinase
ErbB: epidermal growth factor receptor
GEO: Gene Expression Omnibus
Glb1: galactosidase beta 1
H_2_O_2_: hydrogen peroxide
IL-6: interleukin-6
MDM2: murine double minute 2
MI: myocardial infarction
mTOR: mechanistic target of rapamycin
M2: M2-like macrophage
Nrg1: neuregulin 1
Pdgfa: platelet derived growth factor subunit A
PI3K: phosphatidylinositol 3-kinase
qRT-PCR: quantitative reverse transcription-polymerase chain reaction
SA-β-gal: senescence-associated β-galactosidase
SASP: senescence-associated secretory phenotype
Spp1: secreted phosphoprotein 1
Tgfb: transforming growth factor beta
Thy1: thymocyte antigen 1

## Novelty And Significance

### What Is Known?

- M2-like macrophages play a role in post-MI fibrotic tissue formation through activation of cardiac fibroblasts.
- Nrg1/ErbB signaling systems play essential roles in protection and proliferation of cardiomyocytes in response to injury.
- Nrg1/ErbB signaling contributes to macrophage polarization toward an anti-inflammatory phenotype to assist cardiac tissue repair.

### What New Information Does This Article Contribute?

- M2-like macrophage-derived Nrg1/ErbB/PI3K/Akt signaling plays a role in suppressing both senescence and apoptosis of cardiac fibroblasts, which greatly affect fibrotic tissue formation in infarcted adult mouse heart.

M2-like macrophages play an important role in fibrotic tissue formation after myocardial infarction (MI) by activating cardiac fibroblasts. Understanding the effects of apoptosis and senescence on cellular function during tissue repair after MI is important. M2-like macrophages mediate many supplementary benefits in post-MI cardiac repair (e.g., by playing anti-inflammatory and pro-angiogenic roles), in addition to activating cardiac fibroblasts. However, the molecular mechanism by which M2-like macrophages affect the anti-senescence and anti-fibrotic environment after MI has remained unclear. We found that Nrg1, which is known to be expressed by endothelial cells, is also expressed in cardiac CD206^+^F4/80^+^CD11b^+^ M2-like macrophages after MI. In the M2-like macrophage and senescent cardiac fibroblast coculture model, M2-like macrophages suppressed senescence and apoptosis of cardiac fibroblasts, whereas blockade of ErbB function significantly accelerated senescence and apoptosis, resulting in increased collagen synthesis. We showed that the molecular mechanism underlying regulation of fibrotic tissue formation in the infarcted myocardium was in part a reduction in apoptosis and senescence of cardiac fibroblasts by activation of Nrg1/ErbB/PI3K/Akt signaling. Furthermore, systemic blockade of ErbB function exacerbated inflammation and increased accumulation of M2-like macrophages, which resulted in excessive progression of fibrosis. Our finding that Nrg1 critically regulates senescence and apoptosis of cardiac fibroblasts clarifies a part of the mechanism underlying formation of fibrotic tissue after MI. Targeted activation of M2-like macrophages might enhance this endogenous repair mechanism in senescent and apoptotic cardiac fibroblasts and ultimately provide a new treatment option for MI.

## References

1. Zhu F, Li Y, Zhang J, Piao C, Liu T, Li HH and Du J. Senescent cardiac fibroblast is critical for cardiac fibrosis after myocardial infarction. PLoS One. 2013;8:e74535.

2. Dickstein K, Cohen-Solal A, Filippatos G, McMurray JJ, Ponikowski P, Poole-Wilson PA, Stromberg A, van Veldhuisen DJ, Atar D, Hoes AW, Keren A, Mebazaa A, Nieminen M, Priori SG, Swedberg K and Guidelines ESCCfP. ESC Guidelines for the diagnosis and treatment of acute and chronic heart failure 2008: the Task Force for the Diagnosis and Treatment of Acute and Chronic Heart Failure 2008 of the European Society of Cardiology. Developed in collaboration with the Heart Failure Association of the ESC (HFA) and endorsed by the European Society of Intensive Care Medicine (ESICM). Eur Heart J. 2008;29:2388–442.

3. Gemberling M, Karra R, Dickson AL and Poss KD. Nrg1 is an injury-induced cardiomyocyte mitogen for the endogenous heart regeneration program in zebrafish. Elife. 2015;4.

4. Gould KE, Taffet GE, Michael LH, Christie RM, Konkol DL, Pocius JS, Zachariah JP, Chaupin DF, Daniel SL, Sandusky GE, Jr., Hartley CJ and Entman ML. Heart failure and greater infarct expansion in middle-aged mice: a relevant model for postinfarction failure. Am J Physiol Heart Circ Physiol. 2002;282:H615–21.

5. Takemura G, Ohno M, Hayakawa Y, Misao J, Kanoh M, Ohno A, Uno Y, Minatoguchi S, Fujiwara T and Fujiwara H. Role of apoptosis in the disappearance of infiltrated and proliferated interstitial cells after myocardial infarction. Circ Res. 1998;82:1130–8.

6. Shih H, Lee B, Lee RJ and Boyle AJ. The aging heart and post-infarction left ventricular remodeling. J Am Coll Cardiol. 2011;57:9–17.

7. Sharpless NE and Sherr CJ. Forging a signature of in vivo senescence. Nat Rev Cancer. 2015;15:397–408.

8. Munoz-Espin D and Serrano M. Cellular senescence: from physiology to pathology. Nat Rev Mol Cell Biol. 2014;15:482–96.

9. Coppe JP, Desprez PY, Krtolica A and Campisi J. The senescence-associated secretory phenotype: the dark side of tumor suppression. Annu Rev Pathol. 2010;5:99–118.

10. Alam P, Haile B, Arif M, Pandey R, Rokvic M, Nieman M, Maliken BD, Paul A, Wang YG, Sadayappan S, Ahmed RPH and Kanisicak O. Inhibition of Senescence-Associated Genes Rb1 and Meis2 in Adult Cardiomyocytes Results in Cell Cycle Reentry and Cardiac Repair Post-Myocardial Infarction. J Am Heart Assoc. 2019;8:e012089.

11. Hayakawa K, Takemura G, Kanoh M, Li Y, Koda M, Kawase Y, Maruyama R, Okada H, Minatoguchi S, Fujiwara T and Fujiwara H. Inhibition of granulation tissue cell apoptosis during the subacute stage of myocardial infarction improves cardiac remodeling and dysfunction at the chronic stage. Circulation. 2003;108:104–9.

12. Aurora AB, Porrello ER, Tan W, Mahmoud AI, Hill JA, Bassel-Duby R, Sadek HA and Olson EN. Macrophages are required for neonatal heart regeneration. J Clin Invest. 2014;124:1382–92.

13. Shiraishi M, Shintani Y, Shintani Y, Ishida H, Saba R, Yamaguchi A, Adachi H, Yashiro K and Suzuki K. Alternatively activated macrophages determine repair of the infarcted adult murine heart. J Clin Invest. 2016;126:2151–66.

14. Meyer D, Yamaai T, Garratt A, Riethmacher-Sonnenberg E, Kane D, Theill LE and Birchmeier C. Isoform-specific expression and function of neuregulin. Development. 1997;124:3575–86.

15. Fuller SJ, Sivarajah K and Sugden PH. ErbB receptors, their ligands, and the consequences of their activation and inhibition in the myocardium. J Mol Cell Cardiol. 2008;44:831–54.

16. Bersell K, Arab S, Haring B and Kuhn B. Neuregulin1/ErbB4 signaling induces cardiomyocyte proliferation and repair of heart injury. Cell. 2009;138:257–70.

17. Hedhli N, Huang Q, Kalinowski A, Palmeri M, Hu X, Russell RR and Russell KS. Endothelium-derived neuregulin protects the heart against ischemic injury. Circulation. 2011;123:2254–62.

18. Polizzotti BD, Ganapathy B, Walsh S, Choudhury S, Ammanamanchi N, Bennett DG, dos Remedios CG, Haubner BJ, Penninger JM and Kuhn B. Neuregulin stimulation of cardiomyocyte regeneration in mice and human myocardium reveals a therapeutic window. Sci Transl Med. 2015;7:281ra45.

19. Lemmens K, Doggen K and De Keulenaer GW. Role of neuregulin-1/ErbB signaling in cardiovascular physiology and disease: implications for therapy of heart failure. Circulation. 2007;116:954–60.

20. Kirabo A, Ryzhov S, Gupte M, Sengsayadeth S, Gumina RJ, Sawyer DB and Galindo CL. Neuregulin-1beta induces proliferation, survival and paracrine signaling in normal human cardiac ventricular fibroblasts. J Mol Cell Cardiol. 2017;105:59–69.

21. Vermeulen Z, Hervent AS, Dugaucquier L, Vandekerckhove L, Rombouts M, Beyens M, Schrijvers DM, De Meyer GRY, Maudsley S, De Keulenaer GW and Segers VFM. Inhibitory actions of the NRG-1/ErbB4 pathway in macrophages during tissue fibrosis in the heart, skin, and lung. Am J Physiol Heart Circ Physiol. 2017;313:H934–H945.

22. Pascual-Gil S, Abizanda G, Iglesias E, Garbayo E, Prosper F and Blanco-Prieto MJ. NRG1 PLGA MP locally induce macrophage polarisation toward a regenerative phenotype in the heart after acute myocardial infarction. J Drug Target. 2019;27:573–581.

23. Krizhanovsky V, Yon M, Dickins RA, Hearn S, Simon J, Miething C, Yee H, Zender L and Lowe SW. Senescence of activated stellate cells limits liver fibrosis. Cell. 2008;134:657–67.

24. van Deursen JM. The role of senescent cells in ageing. Nature. 2014;509:439–46.

25. Olayioye MA, Neve RM, Lane HA and Hynes NE. The ErbB signaling network: receptor heterodimerization in development and cancer. EMBO J. 2000;19:3159–67.

26. Uray IP, Connelly JH, Thomazy V, Shipley GL, Vaughn WK, Frazier OH, Taegtmeyer H and Davies PJ. Left ventricular unloading alters receptor tyrosine kinase expression in the failing human heart. J Heart Lung Transplant. 2002;21:771–82.

27. Suzuki T, Arumugam P, Sakagami T, Lachmann N, Chalk C, Sallese A, Abe S, Trapnell C, Carey B, Moritz T, Malik P, Lutzko C, Wood RE and Trapnell BC. Pulmonary macrophage transplantation therapy. Nature. 2014;514:450–4.

28. van den Borne SW, Diez J, Blankesteijn WM, Verjans J, Hofstra L and Narula J. Myocardial remodeling after infarction: the role of myofibroblasts. Nat Rev Cardiol. 2010;7:30–7.

29. Shinde AV and Frangogiannis NG. Fibroblasts in myocardial infarction: a role in inflammation and repair. J Mol Cell Cardiol. 2014;70:74–82.

30. Park S, Jiang Z, Mortenson ED, Deng L, Radkevich-Brown O, Yang X, Sattar H, Wang Y, Brown NK, Greene M, Liu Y, Tang J, Wang S and Fu YX. The therapeutic effect of anti-HER2/neu antibody depends on both innate and adaptive immunity. Cancer Cell. 2010;18:160–70.

31. ElZarrad MK, Mukhopadhyay P, Mohan N, Hao E, Dokmanovic M, Hirsch DS, Shen Y, Pacher P and Wu WJ. Trastuzumab alters the expression of genes essential for cardiac function and induces ultrastructural changes of cardiomyocytes in mice. PLoS One. 2013;8:e79543.

32. Bladier C, Wolvetang EJ, Hutchinson P, de Haan JB and Kola I. Response of a primary human fibroblast cell line to H2O2: senescence-like growth arrest or apoptosis? Cell Growth Differ. 1997;8:589–98.

33. Sasaki M, Miyakoshi M, Sato Y and Nakanuma Y. Modulation of the microenvironment by senescent biliary epithelial cells may be involved in the pathogenesis of primary biliary cirrhosis. J Hepatol. 2010;53:318–25.

34. Childs BG, Durik M, Baker DJ and van Deursen JM. Cellular senescence in aging and age-related disease: from mechanisms to therapy. Nat Med. 2015;21:1424–35.

35. Serrano M, Lin AW, McCurrach ME, Beach D and Lowe SW. Oncogenic ras provokes premature cell senescence associated with accumulation of p53 and p16INK4a. Cell. 1997;88:593–602.

36. Kuramochi Y, Cote GM, Guo X, Lebrasseur NK, Cui L, Liao R and Sawyer DB. Cardiac endothelial cells regulate reactive oxygen species-induced cardiomyocyte apoptosis through neuregulin-1beta/erbB4 signaling. J Biol Chem. 2004;279:51141–7.

37. Baliga RR, Pimental DR, Zhao YY, Simmons WW, Marchionni MA, Sawyer DB and Kelly RA. NRG-1-induced cardiomyocyte hypertrophy. Role of PI-3-kinase, p70(S6K), and MEK-MAPK-RSK. Am J Physiol. 1999;277:H2026–37.

38. Deng C, Zhang P, Harper JW, Elledge SJ and Leder P. Mice lacking p21CIP1/WAF1 undergo normal development, but are defective in G1 checkpoint control. Cell. 1995;82:675–84.

39. Wang YA, Elson A and Leder P. Loss of p21 increases sensitivity to ionizing radiation and delays the onset of lymphoma in atm-deficient mice. Proc Natl Acad Sci U S A. 1997;94:14590–5.

40. Alcorta DA, Xiong Y, Phelps D, Hannon G, Beach D and Barrett JC. Involvement of the cyclin-dependent kinase inhibitor p16 (INK4a) in replicative senescence of normal human fibroblasts. Proc Natl Acad Sci U S A. 1996;93:13742–7.

41. Stein GH, Drullinger LF, Soulard A and Dulic V. Differential roles for cyclin-dependent kinase inhibitors p21 and p16 in the mechanisms of senescence and differentiation in human fibroblasts. Mol Cell Biol. 1999;19:2109–17.

42. Yu JS and Cui W. Proliferation, survival and metabolism: the role of PI3K/AKT/mTOR signalling in pluripotency and cell fate determination. Development. 2016;143:3050–60.

43. Ikeda N, Asano K, Kikuchi K, Uchida Y, Ikegami H, Takagi R, Yotsumoto S, Shibuya T, Makino-Okamura C, Fukuyama H, Watanabe T, Ohmuraya M, Araki K, Nishitai G and Tanaka M. Emergence of immunoregulatory Ym1(+)Ly6C(hi) monocytes during recovery phase of tissue injury. Sci Immunol. 2018;3.

44. Acosta JC, Banito A, Wuestefeld T, Georgilis A, Janich P, Morton JP, Athineos D, Kang TW, Lasitschka F, Andrulis M, Pascual G, Morris KJ, Khan S, Jin H, Dharmalingam G, Snijders AP, Carroll T, Capper D, Pritchard C, Inman GJ, Longerich T, Sansom OJ, Benitah SA, Zender L and Gil J. A complex secretory program orchestrated by the inflammasome controls paracrine senescence. Nat Cell Biol. 2013;15:978–90.

45. Tano N, Narita T, Kaneko M, Ikebe C, Coppen SR, Campbell NG, Shiraishi M, Shintani Y and Suzuki K. Epicardial placement of mesenchymal stromal cell-sheets for the treatment of ischemic cardiomyopathy; in vivo proof-of-concept study. Mol Ther. 2014;22:1864–71.

46. Leicht M, Greipel N and Zimmer H. Comitogenic effect of catecholamines on rat cardiac fibroblasts in culture. Cardiovasc Res. 2000;48:274–84.

47. Shintani Y, Kapoor A, Kaneko M, Smolenski RT, D’Acquisto F, Coppen SR, Harada-Shoji N, Lee HJ, Thiemermann C, Takashima S, Yashiro K and Suzuki K. TLR9 mediates cellular protection by modulating energy metabolism in cardiomyocytes and neurons. Proc Natl Acad Sci U S A. 2013;110:5109–14.

