## Supplemental Files for "Nrg1/ErbB Signaling-Mediated Regulation of Fibrosis After Myocardial Infarction"

**Supplementary Materials**

### Supplementary material includes:

Extended Methods

Online Figures I–XII

Online Table I–III

### Extended Methods

#### Animals

Eight- to 10-week-old C57BL/6 mice, purchased from Tokyo Laboratory Animals Science Co., Ltd., were used in the experiments. The mice were maintained in a specific pathogen-free room in our animal facility with a 12-hour light/dark cycle and free access to food and water. *In vitro* and *in vivo* experiments were performed in a blinded manner.

#### Induction of myocardial infarction (MI)

MI was induced in mice by ligating the left coronary artery under 2.0% isoflurane anesthesia and mechanical ventilation, as described previously.[45] Successful establishment of MI was confirmed by changes in the color and motion of the left ventricular wall.

#### In vivo treatment

Mice were injected with 100 µg of trastuzumab (Bio X Cell; catalog #BE0277) on post-MI days 4, 5, and 6. Samples were collected on post-MI days 7, 12, and 28.

#### Isolation of cardiac fibroblasts

Cardiac fibroblasts were isolated from the hearts of Wistar rats (Charles River Laboratories), as described previously[46] but with some modifications. The hearts were cut into 1-mm$^3$ pieces that were plated evenly, without touching each other, in 0.1% gelatin-coated 10-cm dishes. Each fragment was covered by a droplet of Dulbecco's Modified Eagle Medium (DMEM) (Sigma-Aldrich) containing 10% FBS, 50 U/mL penicillin, and 50 µg/mL streptomycin and incubated for

24 hours at 37°C in a $CO_2$ incubator. Sufficient medium was then added to cover the entire bottom of the culture dish, and the culture was incubated for an additional 24 hours. After this period, sufficient medium was added to completely cover the heart fragments, and the culture was continued for an additional 5 days. When fibroblast outgrowth was observed, the cells were collected by trypsinization. The fibroblasts were maintained in DMEM containing 10% FBS, 50 U/mL penicillin, and 50 μg/mL streptomycin in 0.1% gelatin-coated culture flasks and used at passage 3 or 4.

**Preparation of bone marrow-derived macrophages (BMDMs).** Mouse BMDMs were prepared from the femurs and tibiae of 8-week-old wildtype mice, as described previously.[13] Bone marrow mononuclear cells were collected by centrifugation over Ficoll-Paque (GE Healthcare) and were cultivated overnight in a $CO_2$ incubator in DMEM containing 10% fetal bovine serum (FBS), 50 U/mL penicillin, 50 μg/mL streptomycin, and 10 ng/mL granulocyte-macrophage colony stimulating factor (R&D Systems; Catalog #415-ML). Unattached or weakly attached cells were collected and transferred to new dishes and cultivated for an additional 5 days. The cells were then prepared for coculture experiments.

**Coculture of cardiac fibroblasts and BMDMs in a Boyden chamber**

Cardiac fibroblasts ($2 \times 10^4$) were plated on a 0.1% gelatin-coated 6-well dish (Thermo Scientific) and cultured for 48 hours in DMEM containing 10% FBS, 50 U/mL penicillin, and 50 μg/mL streptomycin. The cardiac fibroblasts were subjected to oxidative stress induced with hydrogen peroxide ($H_2O_2$) (Sigma), as described previously[32] but with some modifications. Briefly, cells at passage 3 or 4 were treated with 100 μM $H_2O_2$ for 1 hour and then washed 3 times with phosphate-buffered saline (PBS). The damaged fibroblasts were maintained in DMEM containing 10% FBS, 50 U/mL penicillin, and 50 μg/mL streptomycin for 7 days, and senescence was induced. For coculture experiments, BMDMs ($2 \times 10^4$) were seeded on polycarbonate membrane inserts (pore size: 0.4 μm; Thermo Scientific) that were placed in wells containing senescent fibroblasts. Anti-ErbB4 Ab (10 μg/mL; Thermo Scientific; catalog #MA5-13016) or recombinant Nrg1 (100 ng/mL; MyBioSource; catalog #MBS2009710) was added to the culture medium at the start of the coculture. The senescent fibroblasts and/or BMDMs with or without anti-ErbB4 Ab and with or without recombinant Nrg1 were cultured at 37°C in a humidified $CO_2$ incubator (5% $CO_2$) for 24–48 hours.

**Isolation of heart cells**

Mouse heart cells were isolated as previously described.[47] Mice were killed by cervical dislocation, and immediately thereafter the aorta was clamped and cold Hank's balanced salt solution (HBSS) (Sigma-Aldrich) was injected into the left ventricular cavity. The isolated hearts were cut into 1-$mm^3$ pieces, digested with 0.05% collagenase II (Sigma-Aldrich) at 37°C for 15 minutes and then filtered through a 40-μm cell strainer (BD Falcon). The remnant heart tissues were again digested with fresh digestion solution and similarly filtered. This cycle was repeated 5 times. The suspensions obtained at each cycle were combined and subjected to flow cytometric analyses or fluorescence-activated cell sorting (FACS) after erythrocytes were depleted with Red Cell Lysis Buffer (BioLegend) according to the manufacturer's protocol.

**Flow cytometry and FACS**

The isolated heart cells were resuspended in FACS buffer (HBSS with 2 mM ethylenediaminetetraacetic acid and 0.5% bovine serum albumin) and preincubated with a rat anti-mouse CD16/CD32 Ab at 1:100 dilution (eBioscience; catalog #14-0161) to block Fc receptors. Dead cells and debris were excluded by forward scatter/side scatter and DAPI staining at 1:1,000 dilution (Sigma-Aldrich). To determine phenotypes, the cells were stained with the following for 3 hours at 4 °C: APC-conjugated rat anti-CD11b Ab at 1:100 dilution (eBioscience; catalog #17-0112); phycoerythrin-conjugated rat anti-F4/80 Ab at 1:20 dilution (eBioscience; catalog #12-4801), and Alexa Fluor 488-conjugated rat anti-CD206 Ab at 1:50 dilution (BioLegend; catalog #141709). Cell sorting was performed with a FACSAria II Cell Sorter (BD Biosciences). Cells were acquired from independent biological replicates.

**RNA extraction and real-time polymerase chain reaction (PCR)**

Total RNA was extracted from cells or heart tissue with a Gene Jet PCR purification Kit (Thermo Scientific) and quantified with a Nano-Drop 8000 spectrophotometer (Thermo Scientific). For extraction of total RNA from mouse hearts, fresh tissue was excised from the infarcted area and from a clearly distinguished remote area, and the border area was removed. cDNA was synthesized from 25 ng and 150 ng of total RNA extracted from M2-macrophages and heart tissues, respectively, with a High Capacity cDNA Reverse Transcription Kit (Applied Biosystems). Real-time PCR was performed with QuantStudio (Applied Biosystems) and SYBR Premix Ex Taq II (Takara Bio) under the following conditions: 95°C for 30 seconds followed by 40 cycles at 95°C for 5 seconds and 60°C for 30 seconds. Gene expression levels were normalized to *Gapdh* expression. Gene expression data were acquired from independent biological replicates. The primers are shown in Supplemental Table I.

### Microarray analysis

Total RNA of CD206$^+$F4/80$^+$CD11b$^+$cardiac M2-like macrophages was isolated as described above, amplified with the Ovation RNA Amplification System V2 (NuGEN), and subjected to the Illumina BeadArray platform with a Mouse WG-6 v2.0 Expression BeadChip (Illumina). Two independent biological replicates were prepared for CD206$^+$F4/80$^+$CD11b$^+$ M2-like macrophages, which were isolated from normal hearts and post-MI day 7 hearts. Median per chip normalization was performed for each array. We analyzed only genes for which signal intensity was above 90 in any of the biological duplicates. A 2-fold change cutoff was used to identify differentially expressed genes.

### Immunohistochemistry

Mice were killed by cervical dislocation, and immediately thereafter, the aorta was clamped, and ice-cold PBS was injected into the left ventricular cavity. The heart was then perfused with ice-cold 4% paraformaldehyde in PBS. The heart was removed, cut at the midpoint along the short axis of the left ventricle, embedded in optimal cutting temperature compound (VWR International), and frozen in isopentane chilled in liquid nitrogen. Frozen tissue sections (8 μm thick) were prepared, and non-specific antibody-binding sites were pre-blocked with blocking buffer (PBS containing 5% goat serum). Primary antibodies were then applied overnight at 4°C. After 3 5-minute rinses in PBS, the sections were incubated with appropriate fluorophore-conjugated secondary antibodies and 4′,6-diamidino-2-phenylindole (DAPI) (Sigma-Aldrich; catalog #D9542) in blocking buffer for 1 hour at room temperature. Stained sections were mounted with DAKO Fluorescence Mounting Medium (Agilent, catalog #S302380-2). Data were acquired from independent biological replicates. The primary and secondary antibodies used are shown in Supplemental Table II.

**Picrosirius red staining**

Frozen tissue sections (8 μm thick) were incubated in 1.5% phosphomolybdic acid for 60 minutes, 0.1% picrosirius red solution for 15 minutes, and 0.5% acetic acid solution for 3 minutes. After dehydration in graded ethanol to xylene, the sections were mounted with DPX mounting medium (VWR International). The infarct area was defined as the area with loss of more than 90% cardiomyocytes. Data were acquired from independent biological replicates.

**Masson trichrome staining**

Frozen tissue sections (8 μm thick) were prepared as described above and stained with trichrome (Trichrome Stain Kit, ScyTek Laboratories) according to the manufacturer's instructions. Data were acquired from independent biological replicates.

**Immunocytochemistry**

Cells were fixed in 4% paraformaldehyde/PBS for 5 minutes at room temperature. Cells not set aside for surface antigen staining were incubated in PBS containing 0.1% Triton X-100 for 5 minutes at room temperature. Non-specific antibody binding sites were pre-blocked with PBS containing 5% goat serum for 30 minutes at room temperature. Primary antibodies were then applied to the cells for 1 hour at room temperature: anti-vimentin Ab at 1:100 dilution (Abcam, catalog #24525), anti-cleaved caspase 3 Ab at 1:100 dilution (Cell Signaling; catalog #9661), anti-Ki-67 Ab at 1:100 dilution (eBioscience; catalog #14-5698), and anti-αSMA Ab at 1:00

dilution (Abcam; catalog #5694). Cells were rinsed and then incubated with fluorophore-conjugated secondary antibody at 1:300 dilution (Alexa Fluor 488- or Alexa Fluor 594-conjugated polyclonal antibody, Invitrogen) and DAPI in blocking buffer for 1 hour at room temperature. Data were acquired from independent biological replicates.

**Imaging and analysis**

Digital images were acquired with an all-in-one type fluorescence microscope (BZ-8000; KEYENCE). Images were imported as TIFF files into ImageJ software (National Institute of Health). Images were acquired at 5 separate regions, and the fibrin-positive area of each region was calculated as a percentage of the entire region. The color threshold function of ImageJ was applied to measure macrophages, fibroblasts, the fibrin clot area, and size of the cultured fibroblasts.

**Detection and analysis of proteins**

Protein expression was examined by immunoblotting (Western blotting). Tissue was frozen, crushed, and then lysed in T-PER Tissue Protein Extraction Regent (Thermo Scientific; catalog #78510) according to the manufacturer’s protocol. The protein concentration was measured with the Nano-Drop 8000 spectrophotometer, and lysates containing 50 μg protein each were separated by PAGE, transferred onto a polyvinylidene difluoride membrane, and specific proteins were identified by means of immunoblotting performed with the primary and secondary antibodies shown in Supplemental Table III. Protein bands were visualized with Super Signal West Pico Chemiluminescent Substrate (Thermo Scientific; catalog #34077) according to the manufacturer's instructions. Data were acquired from independent biological replicates.

**Statistics**

All statistical tests were performed with GraphPad Prism version 8 (GraphPad Software). Values are shown mean±SEM. For comparison between multiple sample sets, repeated-measures analysis of variance (ANOVA) or 1- or 2-way ANOVA was performed, followed by Bonferroni's post-hoc test. For comparison between 2 sample sets, 2-tailed, unpaired Student's *t*-test was used. Survival curves were drawn by the Kaplan-Meier method and then compared by log-rank test.

**Online Figure I.**

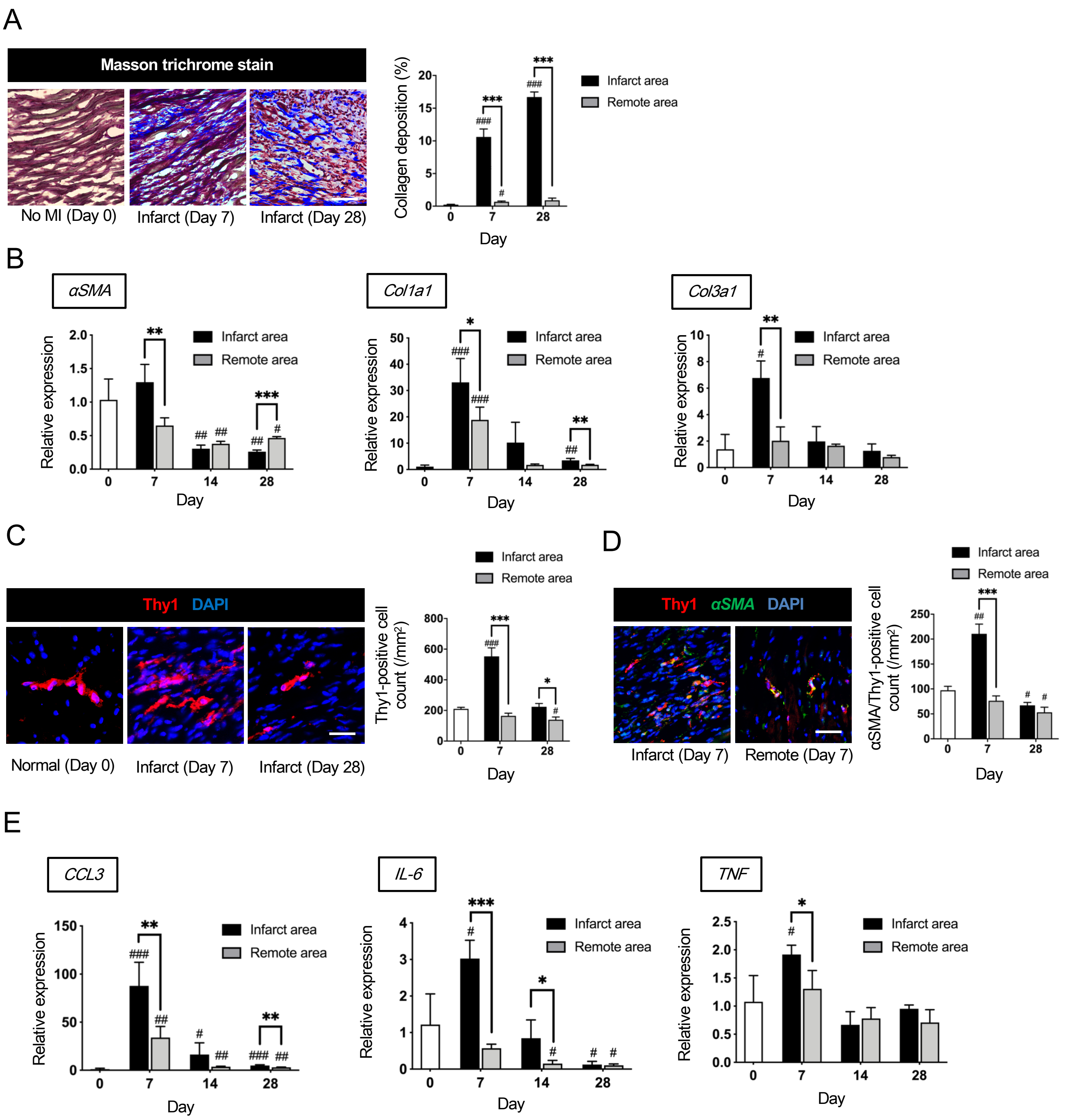


**Online Figure I. Myocardial infarction (MI) promotes activation of cardiac fibroblasts and exacerbates inflammation.**
**A**, Masson trichrome staining revealed increased deposition of collagen fibrils in the infarct area over time after MI. Scale bars: 100 µm. *n*=4 samples each.
**B**, Quantitative reverse transcription-polymerase chain reaction (qRT-PCR) showed post-MI upregulation of fibrosis-associated genes in the infarct area on post-MI day 7, compared with that in the remote area. *n*=4 samples each.
**C**, Immunohistochemistry showed increased accumulation of Thy1$^{+}$ fibroblasts in the infarct area in comparison to that in the remote area. Scale bars: 100 µm. *n*=4 samples each.
**D**, The number of activated cardiac fibroblasts (Thy1$^{+}$ and αSMA$^{+}$ fibroblasts) was significantly increased in the infarct area on post-MI day 7 compared with the number in the remote area. Scale bars: 100 µm. *n*=4 samples each.
**E**, qRT-PCR showed post-MI upregulation of inflammatory genes in the infarct area on post-MI day 7 compared with expression in the remote area. *n*=4 samples each. Gene expression levels relative to those in the non-MI heart (day 0) are shown. Mean±SEM values are shown. **P*<0.05, ***P*<0.01, ****P*<0.005 versus the remote area; $^{\#}P<0.05$, $^{\#\#}P<0.01$, $^{\#\#\#}P<0.005$ versus the non-MI heart; 1-way ANOVA.

### Online Figure II.

A

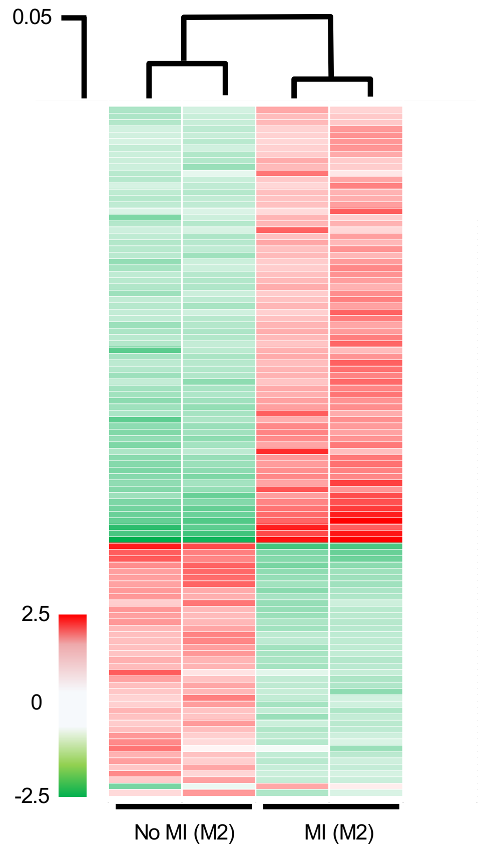


B

| Gene Symbol | Signal intensity | | Rate of change |
|---|---|---|---|
| | no-MI | MI | |
| *F10* | 104.882 | 823.5787 | 7.852431304 |
| *Spp1* | 97.37634 | 388.4397 | 3.989056274 |
| *Inhba* | 90.25688 | 258.7291 | 2.866585905 |
| *Mif* | 413.0435 | 1182.225 | 2.862228797 |
| *Vegfa* | 80.99078 | 218.8935 | 2.702696529 |
| *Slpi* | 82.30864 | 218.8316 | 2.65867107 |
| *C3* | 93.01056 | 230.3347 | 2.476436009 |
| *Nrg1* | 83.75011 | 201.1356 | 2.40161595 |
| *Cxcl9* | 99.36134 | 220.9925 | 2.224129626 |
| *Fn1* | 96.68272 | 199.992 | 2.068539238 |
| *Il1a* | 285.4815 | 574.4752 | 2.012302724 |
| *Il1rn* | 156.1226 | 313.1538 | 2.005819785 |
| *Il10* | 422.77505 | 830.7363 | 1.964960562 |
| *Matn4* | 270.523 | 119.5127 | 0.441783878 |
| *Cyr61* | 243.8087 | 107.6115 | 0.441376784 |
| *Serpinf1* | 657.8257 | 159.2745 | 0.242122647 |

C

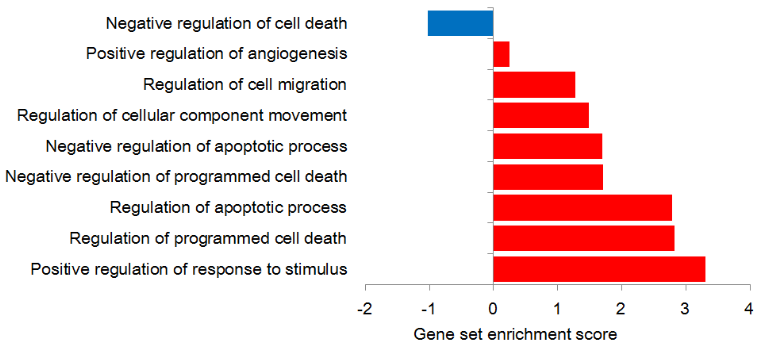


**Online Figure II. Cardiac M2-like macrophages strengthen their reparative ability.**
**A**, $CD206^+F4/80^+CD11b^+$ M2-like macrophages were isolated from normal hearts and from infarcted hearts on day 7 by fluorescence-activated cell sorting (FACS) and subjected to microarray analysis. Macrophages of the different origins showed distinct molecular signatures.
**B**, On FACS, 13 genes encoding secreted proteins were found to be upregulated and 3 were found to be downregulated in M2-like macrophages isolated from infarcted hearts. The upregulated genes included anti-inflammatory and anti-apoptotic genes as well as genes associated with cell survival and tissue repair.
**C**, Gene set enrichment analysis showed that $CD206^+F4/80^+CD11b^+$ M2-like macrophages in the post-MI heart were significantly related to regulation of cell survival and death.

**Online Figure III.**

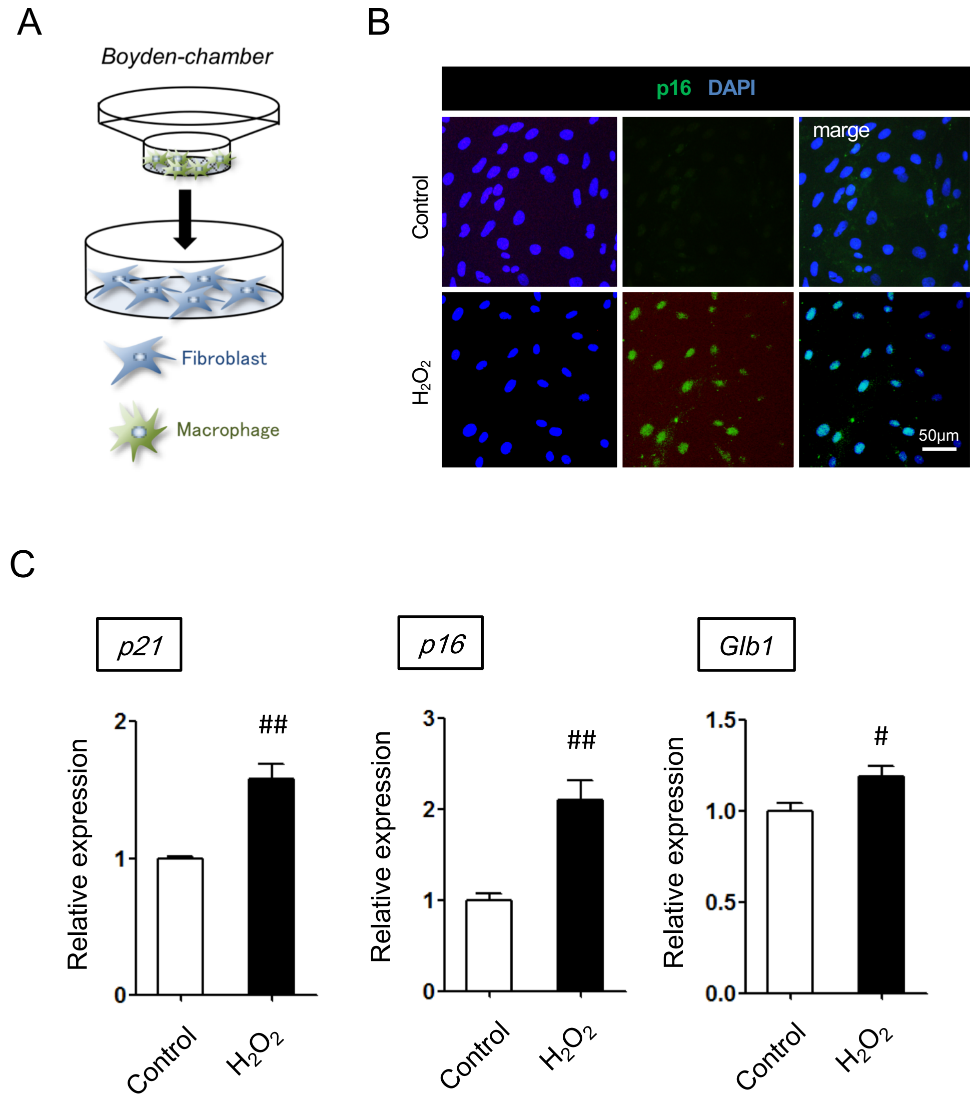


**Online Figure III. Hydrogen peroxide ($H_2O_2$) induces apoptosis and senescence of cardiac fibroblasts.**
**A**, Schematic of the coculture protocol. $H_2O_2$-treated cardiac fibroblasts were cocultured with or without bone marrow-derived macrophages (BMDMs) in a Boyden chamber culture system. Anti-ErbB antibody (Ab) and/or recombinant neuregulin 1 (Nrg1) were added to the relevant samples. The anti-ErbB Ab and/or Nrg1 were added at the beginning of the coculture with BMDMs.
**B**, Representative images of immunocytochemical staining for p16. Seven days after induction of senescence by $H_2O_2$, cultured fibroblasts expressed p16. Nuclei were counterstained with 4′,6-diamidino-2-phenylindole (DAPI). Scale bars: 50 μm.
**C**, Quantitative reverse transcription-polymerase chain reaction analysis confirmed increases in expression of *p21*, *p16*, and *Glb1* in $H_2O_2$-treated fibroblasts. $n$=4 samples each. Gene expression levels relative to the normal cardiac fibroblasts (control) are given. Mean±SEM values are shown.
$^{\#}P<0.05$, $^{\#\#}P<0.01$, $^{\#\#\#}P<0.005$ versus control; by 2-tailed, unpaired Student's $t$-test.

**Online Figure IV.**

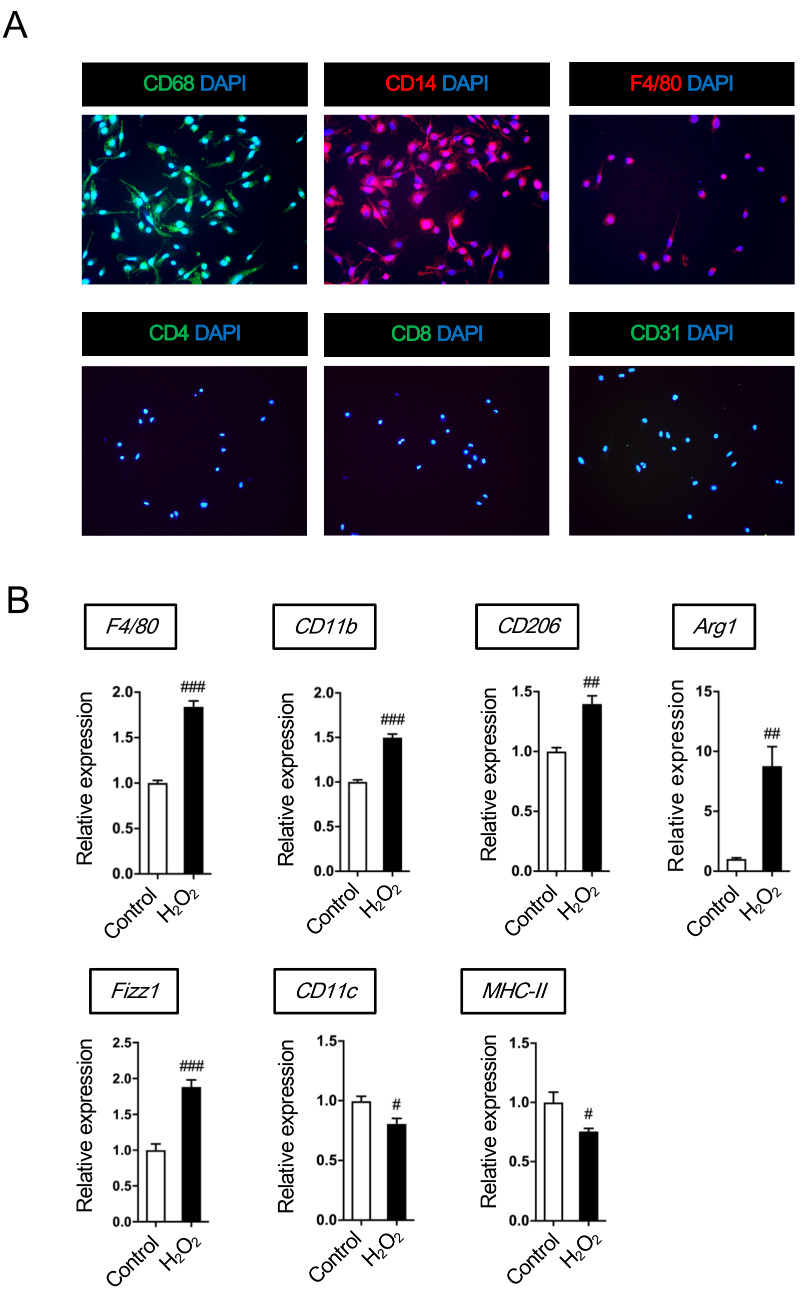


**Online Figure IV. Bone marrow-derived macrophages (BMDMs) cocultured with hydrogen peroxide ($H_2O_2$)-treated cardiac fibroblasts exhibit an M2-like macrophage phenotype.**

**A**, Representative images of immunocytochemical staining for CD68, CD14, F4/80, CD4, CD8, and CD31 in BMDMs. Nuclei were counterstained with 4′,6-diamidino-2-phenylindole (DAPI). Scale bars: 50 µm.

**B**, Quantitative reverse transcription-polymerase chain reaction confirmed increases in expression of pan-macrophage marker genes (*F4/80* and *CD11b*) and M2-like macrophage marker genes (*CD206*, *Arg1*, and *Fizz1*) in BMDMs cocultured with $H_2O_2$-treated fibroblasts. Conversely, expression of M1 macrophage markers (*CD11c* and *MHC-II*) was decreased. *n*=4 samples each. Gene expression levels relative to the BMDMs before coculture (control) are given. Mean±SEM values are shown.

$^{\#}P<0.05$, $^{\#\#}P<0.01$, $^{\#\#\#}P<0.005$ versus control; 2-tailed, unpaired Student's *t*-test.

**Online Figure V.**

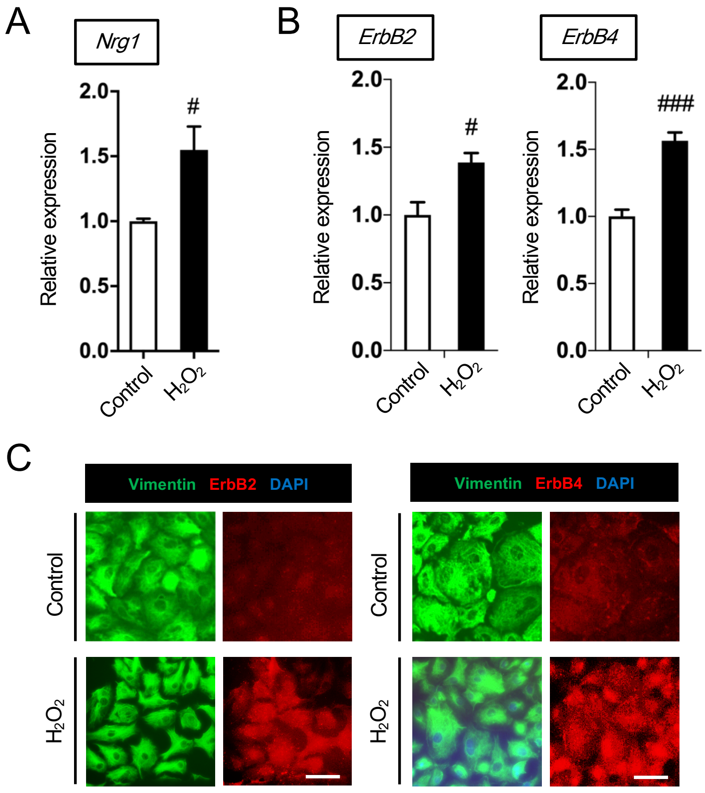


**Online Figure V. Nrg1 expression upregulates in bone marrow-derived macrophages (BMDMs), while cultured fibroblasts express Nrg1 receptors.**

**A**, Quantitative reverse transcription-polymerase chain reaction (qRT-PCR) analysis confirmed an increase in expression of *Nrg1* in BMDMs cocultured with $H_2O_2$-treated fibroblasts compared with that in BMDMs before coculture (control). *n*=4 samples each.

**B**, qRT-PCR analysis confirmed increases in expression of *ErbB2* and *ErbB4* in $H_2O_2$-treated fibroblasts compared with expression in normal fibroblasts (control). *n*=4 samples each.

**C**, Double immunofluorescence staining showed that treatment with $H_2O_2$ enhanced expression of both ErbB2 and ErbB4 receptors on cultured fibroblasts. Nuclei were counterstained with DAPI. Scale bars: 10 µm. Gene expression levels relative to control levels are given. Mean±SEM values are shown.

$^{\#}P<0.05$, $^{\#\#}P<0.01$, $^{\#\#\#}P<0.005$ versus control; 2-tailed, unpaired Student's *t*-test.

**Online Figure VI.**

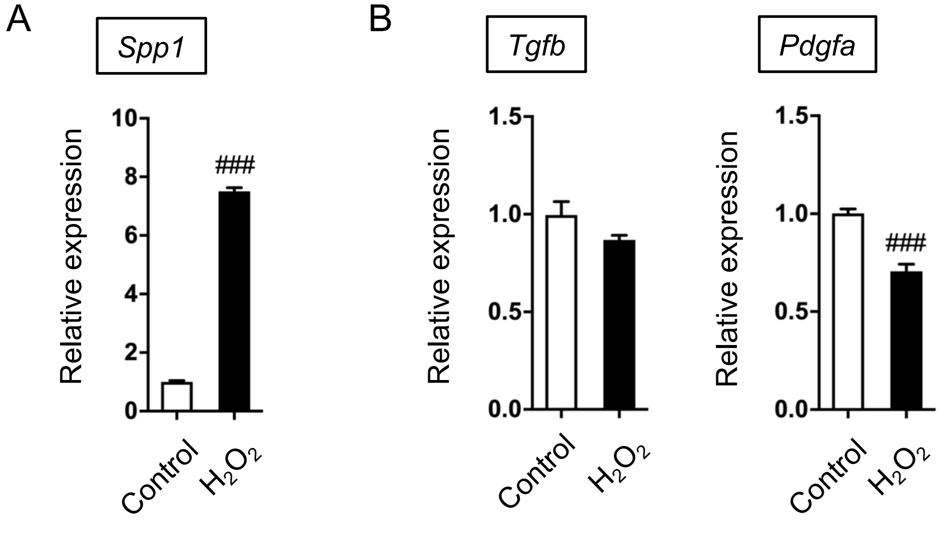


**Online Figure VI. Expression of fibroblast activation-associated gene *Spp1* is increased in bone marrow-derived macrophages (BMDMs).**

**A** and **B**, Quantitative reverse transcription-polymerase chain reaction analysis confirmed that expression of a fibroblast activation-associated gene (osteopontin [*Spp1*]) (A) was increased in BMDMs cocultured with $H_2O_2$-treated fibroblasts. Conversely, expression of other known fibroblast activation-associated genes (transforming growth factor-beta [*Tgfb*] and platelet-derived growth factor subunit A [Pdgfa]) (B) did not increase. $n$=4 samples each. Gene expression levels relative to levels in the BMDMs cocultured with normal fibroblasts (control) are given. Mean±SEM values are shown. $^{\#}P$<0.05, $^{\#\#}P$<0.01, $^{\#\#\#}P$<0.005 versus control; 2-tailed, unpaired Student's *t*-test.

**Online Figure VII.**

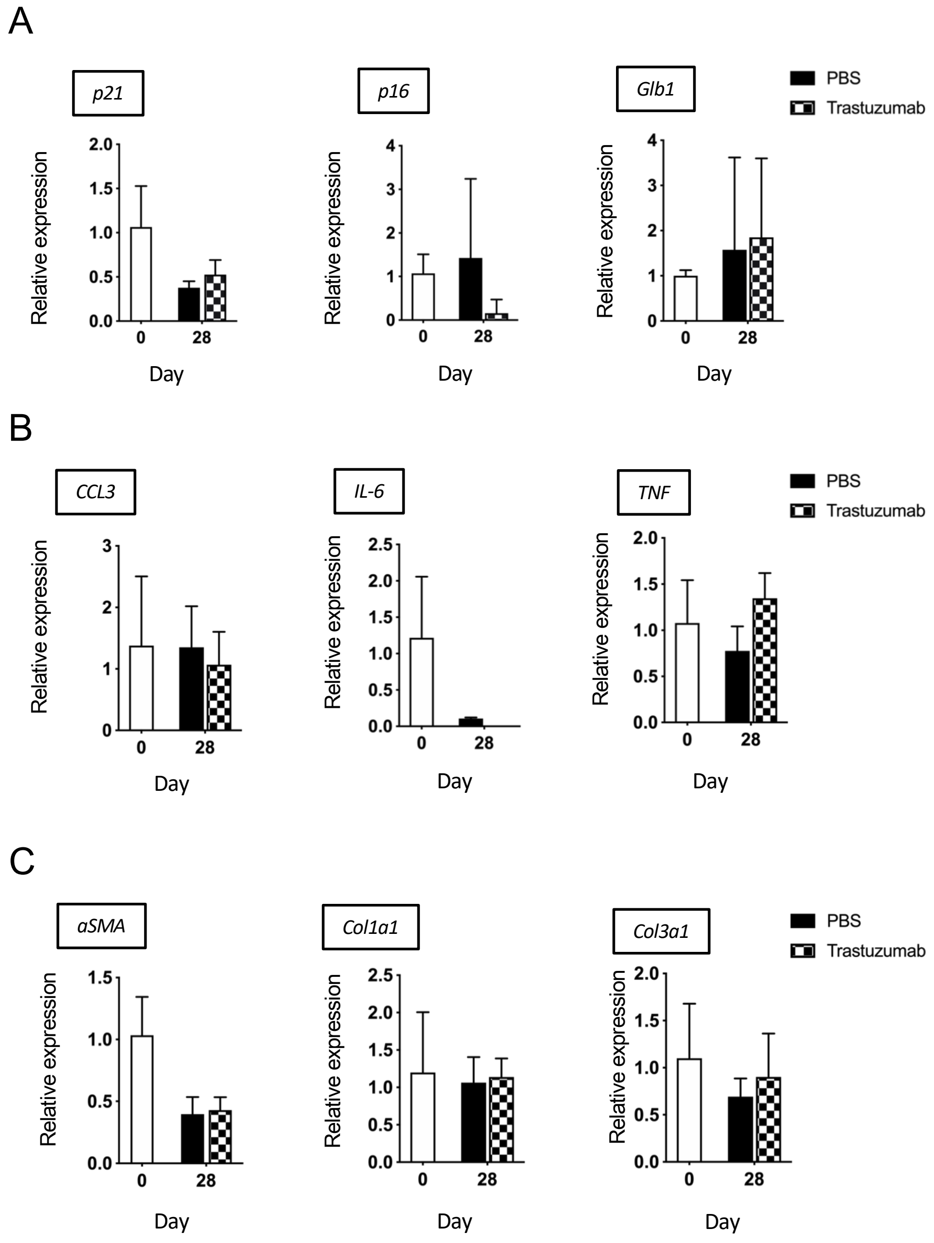


**Online Figure VII. Trastuzumab does not affect apoptosis, senescence, inflammation, or fibroblast activation in the normal heart.**

**A**–**C**, Quantitative reverse transcription-polymerase chain reaction showed no change in mRNA expression levels of (A) senescence-associated genes (*p21*, *p16*, and *Glb1*), (B) senescence-associated secretory phenotype-associated genes (*CCL3*, *IL-6*, and *TNF*), or (C) fibroblast activation-associated gene (*αSMA*) and fibrosis-associated genes (*Col1a1* and *Col3a1*) in mice after intraperitoneal trastuzumab injection. $n$=4 samples each. Gene expression levels relative to those in the normal heart (day 0) are given. Mean±SEM values are shown. *$P$<0.05, **$P$<0.01, ***$P$<0.005 versus sample(s); #$P$<0.05, ##$P$<0.01, ###$P$<0.005 versus the non-MI heart; 1-way ANOVA.

**Online Figure VIII.**

A

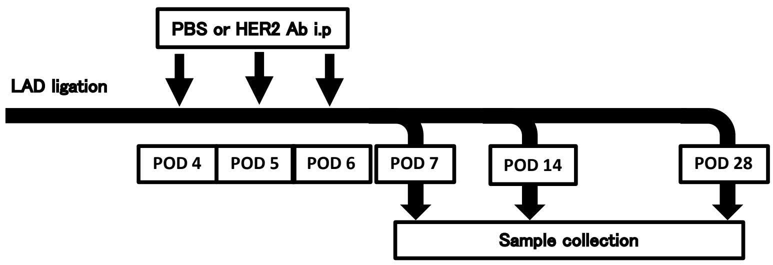


B

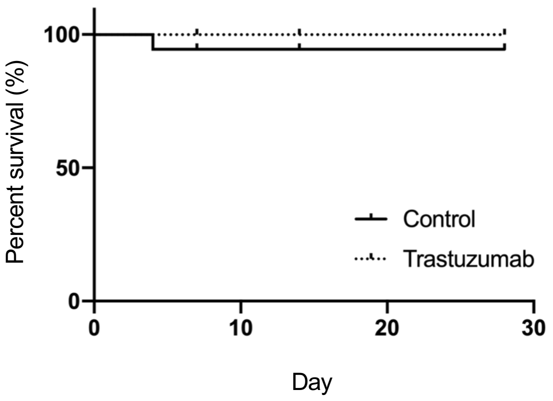


C

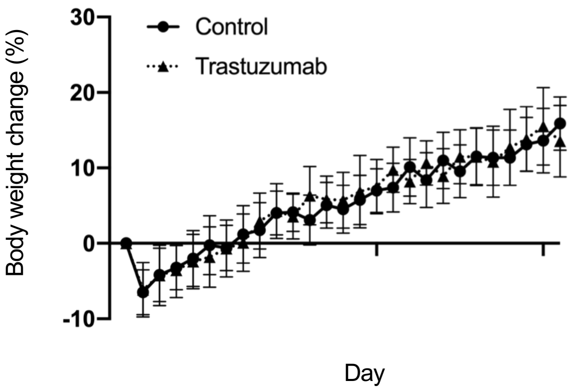


**Online Figure VIII. Trastuzumab does not affect survival or body weight after myocardial infarction (MI)**
**A**, Trastuzumab was injected on post-MI days 4, 5, and 6. Samples were collected on the post-MI days 7, 14, and 28.
**B** and **C**, Intraperitoneal trastuzumab injection did not affect (B) survival or (C) body weight of mice subjected to MI. *n*=12 samples each.

**Online Figure IX.**

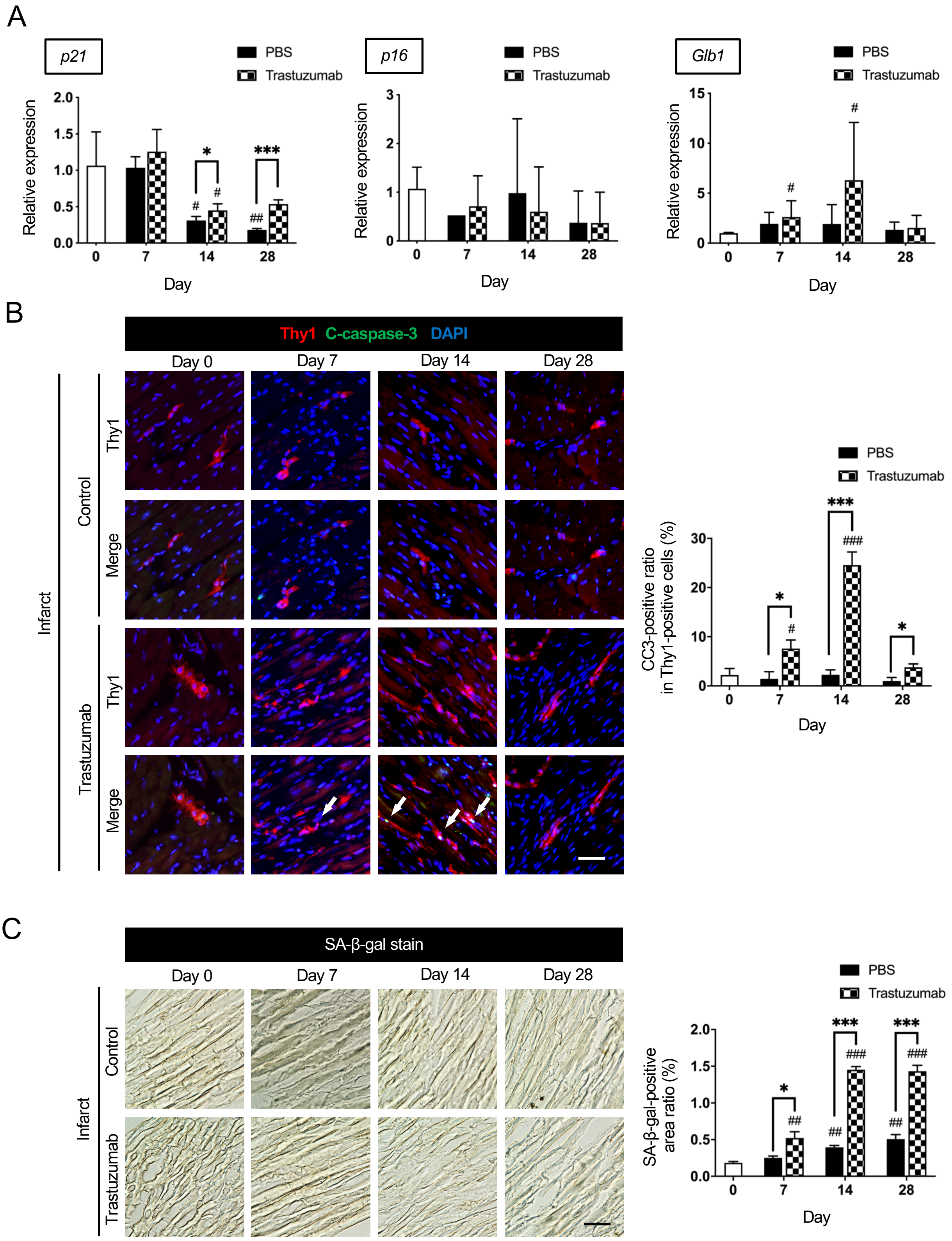

**Online Figure IX. In vivo inhibition of Nrg1 signaling promotes apoptosis and senescence of cardiac fibroblasts even in the remote area.**

**A**, Quantitative reverse transcription-polymerase chain reaction analysis of samples injected with trastuzumab showed upregulation of senescence-associated genes (*Glb1*) in the remote area on post-myocardial infarction (MI) day 14 relatives to expression in vehicle injected post-MI hearts. $n$=4 samples each.

**B**, Double immunofluorescence staining of Thy1 and cleaved caspase 3 (CC3) in samples from mice injected with trastuzumab revealed an increased number of apoptotic cardiac fibroblasts in the remote area on post-MI days 7, 14, and 28 compared with the number in samples from mice injected with vehicle. Arrow shows $Thy1^{+}CC3^{+}$ cells. Scale bars: 100 µm. $n$=4 samples each.

**C**, Staining for senescence-associated beta-galactosidase (SA-β-gal) revealed that, after trastuzumab injection, senescence of cardiac cells in the remote area was significantly exacerbated on post-MI days 7, 14, and 28 compared with that of vehicle-injected post-MI hearts. Scale bars: 100 µm. $n$=4 samples each. Gene expression levels relative to the non-MI heart (day 0) are given. Mean±SEM values are shown.

*$P$<0.05, **$P$<0.01, ***$P$<0.005 versus other sample(s); $^{\#}P$<0.05, $^{\#\#}P$<0.01, $^{\#\#\#}P$<0.005 versus the non-MI heart; 1-way ANOVA.

**Online Figure X.**

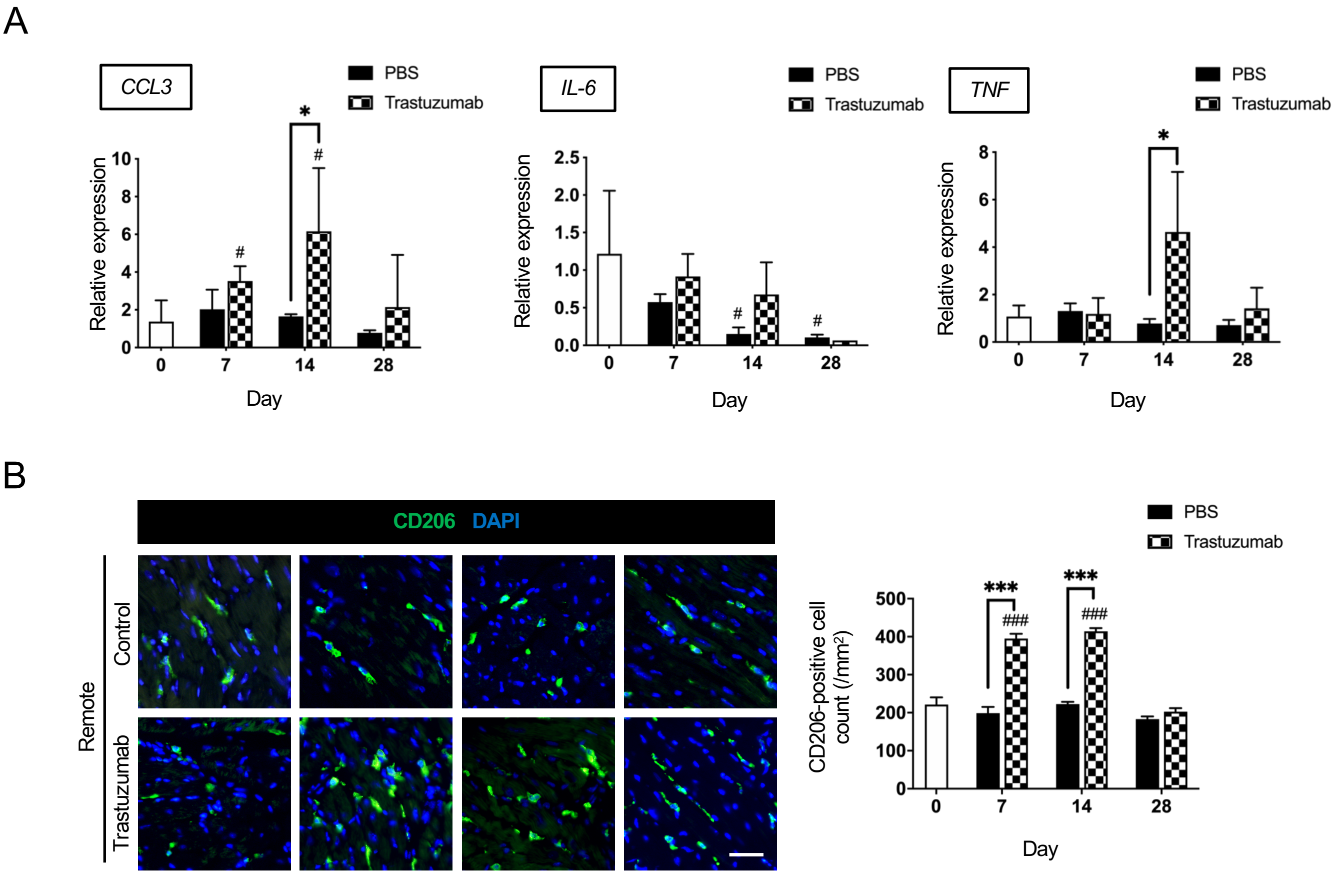


**Online Figure X. In vivo inhibition of Nrg1 signaling exacerbates myocardial inflammation and promotes accumulation of M2-like macrophages even in the remote area**

**A**, Quantitative reverse transcription-polymerase chain reaction analysis of samples obtained from mice injected with trastuzumab showed upregulation of senescence-associated secretory phenotype-associated genes (*CCL3* and *TNF*) in the remote area on post-myocardial infarction (MI) day 14 compared with expression seen in mice injected with vehicle. *n*=4 samples each.

**B**, Immunohistochemistry showed that intraperitoneal injection of trastuzumab significantly accelerated accumulation of CD206$^{+}$ M2-like macrophages in the remote area with a peak on post-MI day 14. Scale bars: 100 μm. n=4 samples each. Gene expression levels relative to those in non-MI heart (day 0) are given. Mean±SEM values are shown. **P*<0.05, ***P*<0.01, ****P*<0.005 versus other samples(s); $^{\#}P$<0.05, $^{\#\#}P$<0.01, $^{\#\#\#}P$<0.005 versus the non-MI heart; 1-way ANOVA.

**Online Figure XI.**

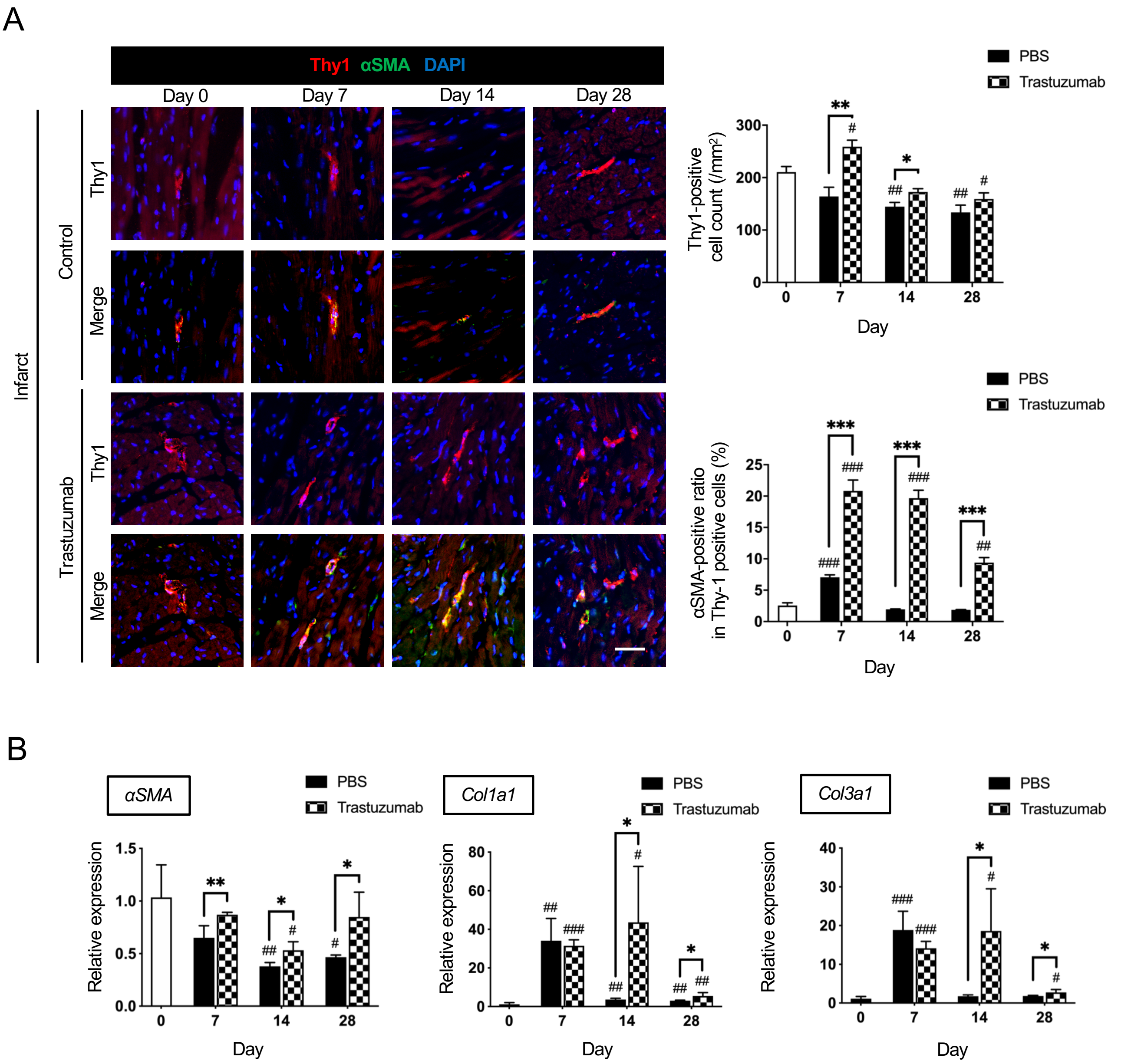


**Online Figure XI.** ***In vivo inhibition of Nrg1 signaling activates cardiac fibroblasts even in the remote area.***
**A**, Double immunofluorescence staining of Thy1 and alpha-smooth muscle actin (αSMA) showed increases in accumulation and activation of cardiac fibroblasts in the remote area of the trastuzumab-injected mice compared with that in the non-trastuzumab-injected (control) mice. Scale bars: 100 µm. *n*=4 samples each.
**B**, Quantitative reverse transcription-polymerase chain reaction analysis showed post-myocardial infarction (MI) upregulation of fibrosis-associated genes (*Col1a1* and *Col3a1*) in the remote area after intraperitoneal trastuzumab injection compared with that in the non-trastuzumab-injected (control) mice on post-MI days 14. *n*=4 samples each. Gene expression levels relative to the non-MI heart (day 0) are given. Mean±SEM values are shown. *$P$<0.05, **$P$<0.01, ***$P$<0.005 versus other sample(s); $^{\#}P$<0.05 $^{\#\#}P$<0.01, $^{\#\#\#}P$<0.005 versus non-MI heart; 1-way ANOVA.

**Online Figure XII.**

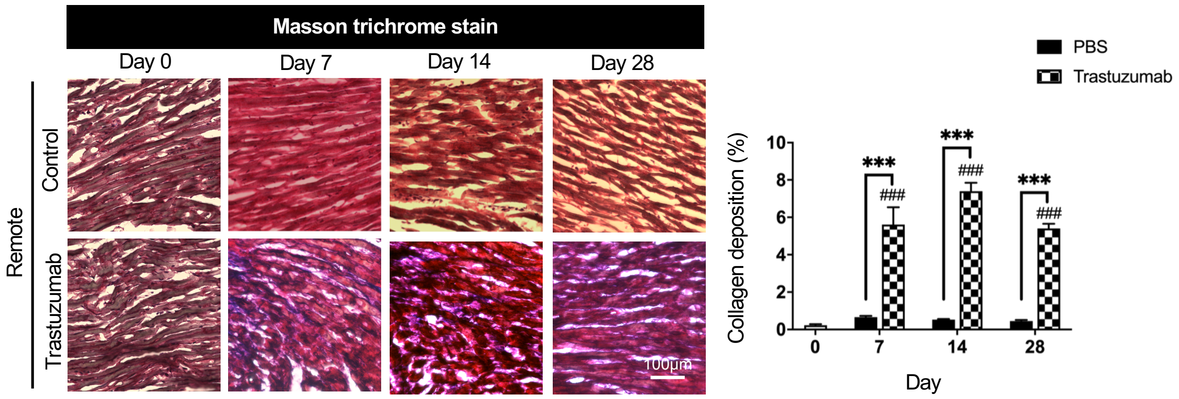


**Online Figure XII.** ***In vivo inhibition of Nrg1 signaling exacerbates fibrosis even in the remote area.***
Masson trichrome staining showed that intraperitoneal trastuzumab injection significantly increased collagen fibrils in the remote area. Scale bars: 100 µm. *n*=4 samples each. Mean±SEM values are shown. *$P$ <0.05, **$P$<0.01, ***$P$<0.005 versus other sample(s); #$P$<0.05 ##$P$<0.01, ###$P$<0.005 versus non-MI heart; 1-way ANOVA.

**Table I. Primers used in PCR**

| | Forward | Reverse |
|---|---|---|
| ***Arg1*** | 5'-CAAGACAGGGCTCCTTTCAG-3' | 5'-AAGCAAGCCAAGGTTAAAGC-3' |
| ***CCL3*** | 5'-CATGAAGGTCTCCACCACTG-3' | 5'-CTCCATATGGCGCTGAGAA-3' |
| ***CD11b*** | 5'-CTGAGAAATGACGGTGAGGA-3' | 5'-CAGCAGGCTTTACAAACCAA-3' |
| ***CD11c*** | 5'-GGTCCTACTGTGCACCACAC-3' | 5'-GACACTCCTGCTGTGCAGTT-3' |
| ***CD206*** | 5'-TGATTACGAGCAGTGGAAGC-3' | 5'-GTTCACCGTAAGCCCAATTT-3' |
| ***p21 (Cdkn1a)*** | 5'-GATCCACAGCGATATCCAGACA-3' | 5'-AGAGACAACGGCACACTTTG-3' |
| ***p16 (Cdkn2a)*** | 5'-ATAGACTAGCCAGGGCAGCG-3' | 5'-TTGCCCATCATCATCACCTGT-3' |
| ***Col1a1*** | 5'-TGAGCCAGCAGATTGAGAAC-3' | 5'-CCAGTACTCTCCGCTCTTCC-3' |
| ***Col3a1*** | 5'-AGTCTGGAGTCGGAGGAATG-3' | 5'-AGGATGTCCAGAGGAACCAG-3' |
| ***ErbB2*** | 5'-CTGAATACCATGCAGATGGG-3' | 5'-TCACACCATAGCTCCACACA-3' |
| ***ErbB4*** | 5'-GAGGAAAGCCCTATGATGGA-3' | 5'-TCCAACATTTGACCATGACC-3' |
| ***F4/80*** | 5'-CAACCTGCCACAACACTCTC-3' | 5'-CCACATCTTCACAGGATTCG-3' |
| ***Fizz1*** | 5'-AGGAACTTCTTGCCAATCCA-3' | 5'-ACAAGCACACCCAGTAGCAG-3' |
| ***Glb1*** | 5'-CGCTACATCTCGGGAAGCAT-3' | 5'-GGGCACGTACGTCTGGATTG-3' |
| ***Gapdh*** | 5'-CCCCTGGCCAAGGTCATCCA-3' | 5'-CGGAAGGCCATGCCAGTGAG-3' |
| ***IL-1α*** | 5'-CCATGATCTGGAAGAGACCA-3' | 5'-GACAAACTTCTGCCTGACGA-3' |
| ***IL-6*** | 5'-AGTCCGGAGAGGAGACTTCA-3' | 5'-TTGCCATTGCACAACTCTTT-3' |
| ***Ki-67*** | 5'-TATCTGGGCCACCTACCTTC-3' | 5'-GCTGTTTCCAGTCCGCTTAC-3' |
| ***MHC-II*** | 5'-ATTGCGAAAGCTGCAGAAC-3' | 5'-TAGCAGCCAGTCATCCTTTG-3' |
| ***Nrg1*** | 5'-GAATTTATGGAAGCGGAGGA-3' | 5'-CAGTAGGCCACCACACACAT-3' |
| ***Spp1*** | 5'-GAGGAAACCAGCCAAGGTAA-3' | 5'-TAGTCCCTCAGAATTCAGCCA-3' |
| ***Pdgfα*** | 5'-GAGGAGGAGACAGATGTGAGG-3' | 5'-ATTGGCAATGAAGCACCATA-3' |
| ***Tgfβ*** | 5'-CAACTTCTGTCTGGGACCCT-3' | 5'-CGGGTTGTGTTGGTTGTAGA-3' |
| ***Tnf*** | 5'-TCGTAGCAAACCACCAAGTG-3' | 5'-TTGTCTTTGAGATCCATGCC-3' |
| ***p53(Trp53)*** | 5'-GTTCCGAGAGCTGAATGAGG-3' | 5'-TTATGGCGGGAGGTAGACTG-3' |
| ***Vegfr2*** | 5'-TGTGGCTTCCTGATGGCAGAA-3' | 5'-AGAAACCAGTAGACATAGTTT-3' |
| ***Vegfα*** | 5'-GTACCTCCACCATGCCAAGT-3' | 5'-GCATTCACATCTGCTGTGCT-3' |

**Table II. Antibodies used for immunocytochemistry**

| Primary antibodies | Host | Dilution | Company / Catalogue No. |
|---|---|---|---|
| **AlexaFluor488-conjugated anti-CD206** | Rat | 1:100 | BioLegend / 141709 |
| **anti-ErbB2** | Rabbit | 1:100 | Abcam / Ab214275 |
| **anti-ErbB4** | Hamster | 1:100 | AbD Serotec / MCA1369 |
| **anti-CD90 (Thy1)** | Rat | 1:100 | eBioscience / 14-0901 |
| **anti-αSMA** | Rabbit | 1:100 | Abcam / ab5694 |
| **anti-Ki67** | Rat | 1:100 | eBioscience / 14-5698 |
| **anti-Cleaved caspase-3** | Rabbit | 1:200 | Cell Signaling / 9661 |
| **anti-Vimentin** | Chicken | 1:100 | Abcam / ab24525 |
| **Secondary antibodies** | | | |
| **AlexaFluor 488-conjugated antibody** | Goat | 1:300 | Invitrogen / A-11006 |
| **AlexaFluor 488-conjugated antibody** | Donkey | 1:300 | Invitrogen / A-21208 |
| **AlexaFluor 488-conjugated antibody** | Goat | 1:300 | Invitrogen / A-11039 |
| **AlexaFluor 594-conjugated antibody** | Goat | 1:300 | Invitrogen / A-11007 |
| **AlexaFluor 594-conjugated antibody** | Goat | 1:300 | Invitrogen / A-11012 |

**Table III. Antibodies used for immunoblotting**

| Primary antibodies | Host | Dilution | Company / Catalogue No. |
|---|---|---|---|
| **Anti-PI3K p85 alpha (phospho Y607)** | Rabbit | 1:1000 | Abcam / ab182651 |
| **Anti-phospho-Akt (Ser473)** | Rabbit | 1:2000 | Cell Signaling / 4060 |
| **HRP-conjugated anti-PI3K** | Rabbit | 1:5000 | Abcam / ab200773 |
| **HRP-conjugated anti-Akt** | Rabbit | 1:1000 | Cell Signaling / 8596 |
| **HRP-conjugated anti-beta Actin** | Mouse | 1:50000 | Abcam / ab49900 |
| **Secondary antibody** | | | |
| **HRP-conjugated anti-rabbit IgG** | Goat | 1:1000 | Cell Signaling / 7074S |
